# Dynamic time-locking mechanism in the cortical representation of spoken words

**DOI:** 10.1101/730838

**Authors:** A. Nora, A. Faisal, J. Seol, H. Renvall, E. Formisano, R. Salmelin

**Author notes:** These authors contributed equally to this work (A. Nora, A. Faisal).

## Abstract

Human speech has a unique capacity to carry and communicate rich meanings. However, it is not known how the highly dynamic and variable perceptual signal is mapped to existing linguistic and semantic representations. In this novel approach, we utilized the natural acoustic variability of sounds and mapped them to magnetoencephalography (MEG) data using physiologically-inspired machine-learning models. We aimed at determining how well the models, differing in their representation of temporal information, serve to decode and reconstruct spoken words from MEG recordings in 16 healthy volunteers. We discovered that time-locking of the cortical activation to the unfolding speech input is crucial for the encoding of the acoustic-phonetic features. In contrast, time-locking was not highlighted in cortical processing of non-speech environmental sounds that conveyed the same meanings as the spoken words, including human-made sounds with temporal modulation content similar to speech. The amplitude envelope of the spoken words was particularly well reconstructed based on cortical evoked responses. Our results indicate that speech is encoded cortically with especially high temporal fidelity. This mechanism may contribute to the frequently reported entrainment of the cortical oscillations to the amplitude envelope of speech. Furthermore, the phoneme content was reflected in cortical evoked responses simultaneously with the spectrotemporal features, pointing to an instantaneous transformation of the unfolding acoustic features into linguistic representations during speech processing.

**Summary:** Computational modeling of cortical responses to spoken words highlights the relevance of temporal tracking of spectrotemporal features, which is likely pivotal for transforming the acoustic-phonetic features into linguistic representations.

## Introduction

Humans effortlessly recognize and react to a variety of natural sounds, but are especially tuned to speech, which is a highly effective way to convey meanings. However, it remains unclear whether speech sounds are processed and represented in a unique fashion in the brain. Numerous studies have attempted to localize speech-specific processing stages [1-3], but while subtle differences in the time sequence of activation for processing speech vs. other meaningful sounds have been described [4, 5], it has remained unclear how they relate to possible unique computations in speech processing. A major challenge in understanding speech processing has been to describe, conceptually and mechanistically, how the brain matches highly variable acoustic signal to linguistic representations, when there is no one-to-one correspondence between the two. It has been suggested that intermediate phonetic/phonological representations act as an interface between auditory and lexical levels. Therefore, the detailed analysis of temporal modulations that distinguish between phonemes may be crucial for speech processing [2, 6]. Sensitivity to spectro-temporal features of speech that correspond to different phonemes has been observed in superior temporal cortex [7, 8]. Recently, superior temporal areas have been demonstrated to show sensitivity for the temporal structure of speech sounds but not for other sounds with similar acoustic content [9]. Thus, the key to understanding the mapping of acoustic signal to linguistic representations during speech processing may lie in tracking its temporal properties, and this may be how the processing of speech differs from the processing of other sounds.

The crucial question of whether there is something special in the cortical processing of speech has often been approached through careful acoustic matching of linguistic and nonlinguistic stimuli and comparison of their activation patterns in the brain [9-11]. The meticulous use of acoustically modified sounds has shown that while primary auditory cortices respond identically to speech and acoustically matched non-speech sounds, responses in nonprimary regions differ [9, 12]. However, natural, meaningful and behaviorally relevant sounds might more accurately capture the cortical sensitivity to the most relevant spectrotemporal features for sound processing [13]. Also, to ensure similar attentional engagement and endpoint of processing, the speech and non-speech sounds should refer to corresponding meanings. Therefore, here we take a different approach to uncover the possible speech-specific mechanism of cortical processing. Our stimuli are natural spoken words and environmental sounds and, instead of delimiting their acoustic properties, we make full use of their large natural variability. We model representations of the different sounds employing multiple physiologically-inspired computational models and use advanced machine-learning techniques to test how well each of the models can account for and predict the cortical representations of the spoken words. Each model attempts to use the variation in the MEG response signals to explain variation within each stimulus feature. Thus the ability of each model to predict and reconstruct the sounds reveals whether the underlying assumptions of the model are an accurate description of how human brain encodes the stimulus features. This approach effectively renders a direct comparison of cortical responses to speech versus other sounds unnecessary. The systematic comparison of the same models within spoken words and separately within environmental sounds allows us to draw conclusions on possible differences in cortical encoding of speech versus other sounds.

Previous machine-learning-based neural decoding models [14] combined with models of cortical processing of sounds [15] have demonstrated tonotopy as well as selectivity for specific spectrotemporal modulations in the auditory cortical areas [16], and allowed successful reconstruction of spoken words based on fMRI and intracranial recordings [17-19]. These findings suggest that perceptually relevant features of speech, e.g. slow temporal modulations, are enhanced in population-level cortical processing. Phonemes have been successfully decoded in other studies [7, 8, 20, 21]. On even more abstract levels of representation, while semantic features of visual stimuli have been successfully decoded [14], semantic classification of spoken words has so far been reported only for small sets of stimuli [22-24]. In a different vein of studies, temporal locking (‘entrainment’) of cortical oscillations to the amplitude envelope of speech has emerged as a potential mechanism for mapping between multiscale spectrotemporal features and linguistic structures [25, 26]. The particular importance of the temporal envelope of speech has been demonstrated also for speech recognition [27, 28]. However, temporal tracking of the continuous speech signal also depends on fine spectrotemporal detail [29]. In a recent study, low-frequency cortical oscillatory activation to speech appeared best modeled as a combination of spectrotemporal features and phonetic-feature labels, highlighting the tracking of the spectrotemporal detail within speech [21].

However, whether such temporal tracking is particularly relevant for the processing of speech, and less crucial when processing other types of sounds, remains an open question. It has been well established that subcortical and cortical neurons, starting from the cochlea, show phase-locked encoding of both environmental and speech sounds [30, 31], and neural entrainment is observed also for other sounds than speech, for example music [32]. In contrast, recent studies have suggested that spectrotemporal modulations corresponding to different phonetic features might not be best represented in a linear, time-locked manner [15, 17]. Along these lines, accurate reconstruction of speech can be achieved from fMRI recordings by modeling the encoding of the speech signal as a selection of frequency-specific spectro-temporal modulation filters, despite the inherent loss of temporal information due to the sluggish hemodynamics and poor temporal sampling of the blood oxygen level-dependent (BOLD) response [16, 18]. This suggests spatially distributed representations of temporal information, similarly to frequency content, and questions the crucial role of temporal tracking of the speech signal. To this end, machine learning investigation that combines time-sensitive measures of cortical activation to acoustic models differing in their temporal detail could shed light on the importance of time-locked cortical encoding for the parsing of the phoneme content and online comprehension of individual words.

In this study, we evaluate the hypothesis that temporal tracking of the spectrotemporal content is particularly important in the cortical encoding of spoken words and not as essential for processing of other sounds, by modeling and decoding acoustic representations of the sounds, as well as the phoneme sequence of words. We use multiple acoustic representations of the sounds that differ in temporal detail: time-averaged spectral content, time-averaged spectrotemporal modulation content, time-resolved spectrogram, and amplitude envelope. Cortical evoked activation is measured with MEG, which detects cortex-wide neuronal signaling on a millisecond scale and is thus an efficient tool for studying the temporal properties of sound encoding at different levels of processing. However, an important challenge in predicting the temporal evolution of sound features from high-dimensional time-sensitive MEG recordings has been the lack of rapidly computable and scalable computational models. In the current study, decoding of the time-varying spectrogram, amplitude envelope, and the phoneme sequence of the sounds is achieved using a convolution model [17, 33, 34], with a new formulation that efficiently handles the high spatiotemporal dimensionality of MEG data [35]. The convolution model predicts each time point in the stimulus features (spectrogram / amplitude envelope / phoneme sequence) based on a time window in the unfolding MEG evoked signals, thus assuming that the activation of neuronal populations follows closely in time the time-sequence of stimulus features. Decoding of the time-averaged spectral and spectrotemporal modulation content as well as the semantic features (i.e. non-time-varying features) of the sounds can be done with a regression model [14, 16, 36], where each stimulus feature is predicted based on all time points of the MEG responses, and thus no such time-locking is assumed. The accuracy of these models is evaluated by telling apart two new sounds based on their respective brain activation patterns [36, 37]. Related models have been developed and successfully applied for analysis of multimodal data [21, 38-41]. The essential novel contribution of the present study is the systematic comparison of multiple models that differ in their handling of temporal information to decode sounds and, thereby, to infer the nature of the underlying neural representations.

To evaluate whether the same processing principles apply to other meaningful sounds beyond speech, or not, the same machine learning models are separately employed to decode among a wide range of environmental sounds, within the same individuals. Similar attentional engagement and endpoint of processing for the spoken words and environmental sounds was ensured by choosing sets of items that refer to corresponding meanings (e.g. the uttered word “cat” to study speech encoding, and the sound of a cat “meow” to study environmental sound encoding). The same set of semantic features was used for modeling both classes of sounds to ensure that they equally activated similar meanings. By evaluating the performance of time-varying and non-time-varying machine learning models in decoding natural spoken words and, in parallel, quantifying the relative performance of these same models in decoding natural non-speech sounds, the present study offers essential new insights into cortical mechanisms that are potentially crucial for cortical encoding of speech.

## Results

The speech stimuli were 44 words from various semantic categories, spoken by males, females and children to make them acoustically variable (**Fig. S1**, **Table S1**). Cortical responses were recorded with MEG in 16 healthy adults. Each stimulus was presented 20 times during the experiment. To ensure concentration, participants performed a one-back task where they identified immediate repetitions of the same meaning, i.e., two different speakers speaking the same word or two different exemplars of the same environmental sounds, presented one after the other (**Methods**). In visual inspection, the sounds showed typical auditory evoked responses at the sensor-level (**Fig. S2a**) and, based on source localization by Minimum Norm Estimates (MNEs), prominent activation around the auditory cortices in the left and right temporal lobes (**Fig. S2b**).

### Acoustic decoding of spoken words relies on neural responses that closely follow time-varying spectral features

Each decoding model was trained and tested separately for each individual participant. Prior to performing machine learning analysis, the stimulus features and MEG signals were standardized across all stimuli. Thus, the absolute power did not affect model estimation. Instead, the models used variation in the MEG signal power to predict variation in each stimulus feature, across stimuli. In the training phase, each model learned a mapping between a stimulus feature set and the MEG data based on all but two sounds (**Fig. 1a&b**). The model was then evaluated on the remaining data from the same participant, i.e. used to tell apart the two held-out words/environmental sounds based on their predicted features (leave-two-out cross-validation, **Fig. 1c**; **Methods**). The training and testing steps were repeated for all possible sound pairs.

**Figure 1.**
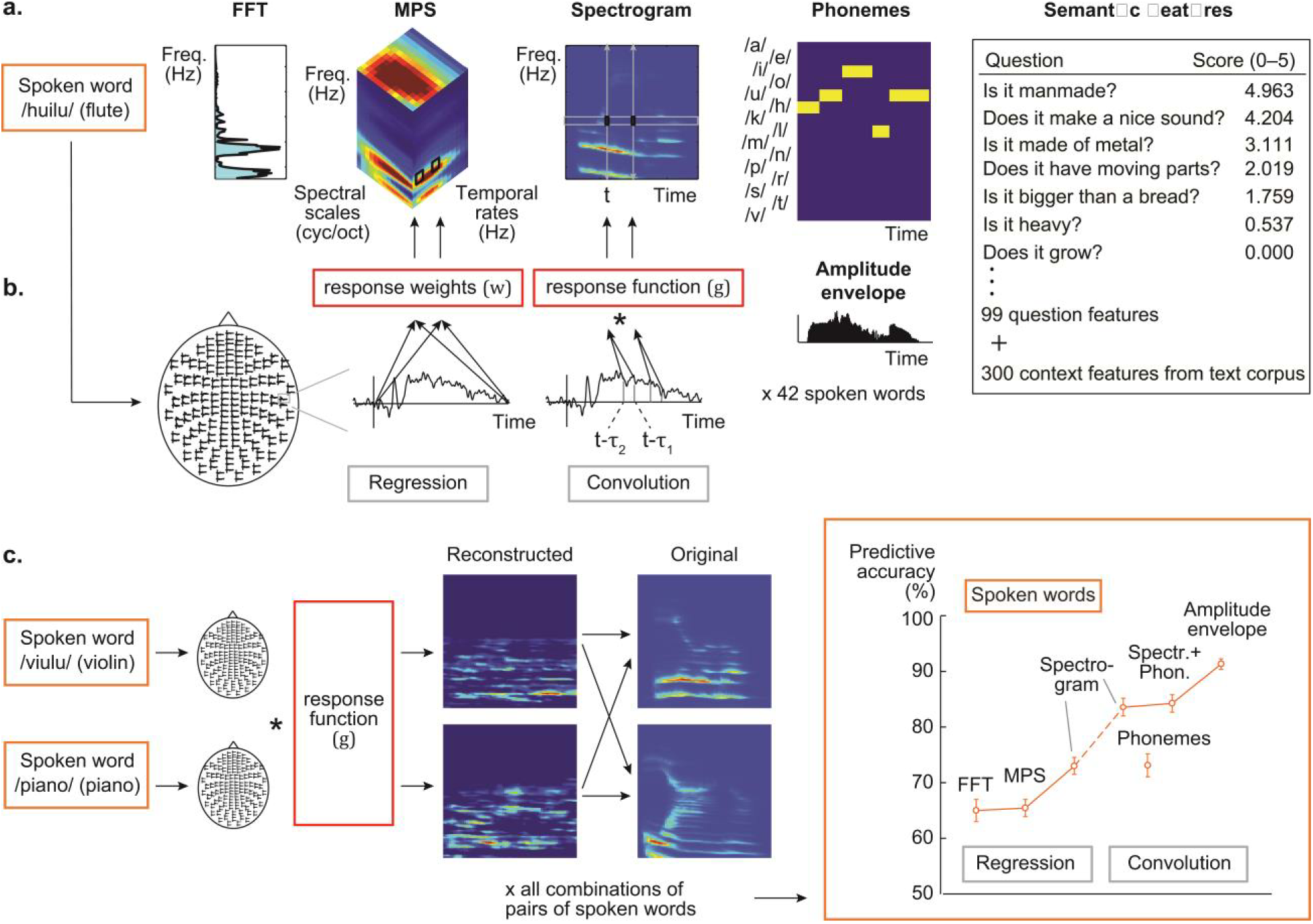
Different decoding models and their prediction accuracy. **a) Stimulus features** for a spoken word. The Fourier transform (FFT) model represents the frequency spectrum of the sound extracted in 128 frequency bands with logarithmically spaced center frequencies (180–7246 Hz). The modulation power spectrum (MPS) model represents energy in four spectral scales (wide to narrow) and four temporal modulation rates (slow to fast), averaged over time. The sound spectrogram quantifies the time-evolving frequency content of the sound extracted in 128 frequency bands and 10-ms time windows by using short-term FFT. The amplitude envelope carries the temporal changes without frequency information (frequency axis averaged out; shown below the phoneme model). The phoneme sequence is the phoneme annotation of the word for each 10-ms time window. Semantic features were represented by scores of 99 questions (a few example questions shown) and a 300-dimensional vector trained with the co-occurrences of context words (words occurring near the stimulus words) in a large text corpus using the word2vec algorithm. **b) Model estimation** for the regression model (left) and the convolution model (right). A mapping between the cortical MEG responses (here illustrated on one sensor) and each stimulus feature was learned with kernel regression or kernel convolution model. The regression weights or response function of the convolution model are learned based on training data (all but two sounds). As illustrated here, the regression model predicts power at each frequency-rate-scale point of the MPS by multiplying cortical responses at all (or selected) time points with unknown weights (*w*). The convolution model predicts the amplitude at each time-frequency point of the spectrogram by convolving the time-sequence of cortical responses with an unknown spatiotemporal response function (*g*). Specifically, values at each frequency band of a new spoken word are predicted at each time point *t* (moving from 0 to end of the sound) based on MEG responses in the time range from (*t − τ_2_*) to (*t − τ_1_*), here illustrated for the lag window *−τ_2_* = 100 to *−τ_1_*= 180 ms at time points *t*. **c) Model testing** aimed to tell apart two left-out sounds by reconstructing the sound features (here, spectrograms) and correlating them with the original spectrograms. The procedure was repeated for all possible pairings of sounds. Predictive accuracy for the spoken words across all test sound pairs and 16 participants (mean ± SEM) is shown on the right. The regression model was used for the decoding of non-time-varying features (FFT frequency bins / MPS rate-scale-frequency points / semantic question scores and corpus statistics) and the convolution model was used for decoding of time-varying features (spectrogram time-frequency points / phonemes at each time point / amplitude envelope); for a control analysis, the spectrogram was also decoded with the regression model. Predictive accuracy improved markedly when spoken words were modeled with the convolution spectrogram model, formalizing the concept that the neuronal population response follows closely in time the unfolding time-sequence of the sound acoustics, i.e. each frequency band of the stimulus spectrogram. Even better performance was obtained with the combination where the spoken words were modeled using both the spectrogram and the phoneme sequence descriptions, and the convolution model linked those descriptions to the MEG signals (significant difference to using the spectrogram description alone, p = 0.01). The best performance in spoken word classification was obtained using the sounds’ amplitude envelope.

As decoding models, we compared four sets of acoustic features that vary with respect to whether and how they incorporate information about the temporal modulations within the spoken words (**Fig. 1a, Methods**). The simplest set of features was based on the fast Fourier transform (*FFT*) which contains no information about the temporal evolution of the sound and represents the frequency content of the sound across its entire duration, calculated with a filter bank of 128 frequency bands.

In addition to displaying an organization by frequency, the primary and secondary auditory cortices respond to different rates of temporal modulations at different spectral scales [16-18]. These intricate multidimensional modulations within sounds are captured by our second set of features, the modulation power spectrum (*MPS*). The MPS (also called modulation transfer function, or MTF, when describing neuronal filter properties [42]) represents energy in wide to narrow spectral scales and at slow to fast temporal modulation rates at different frequency bands (same as in FFT) for the entire duration of the sound [15]. This type of time-averaged modulation content has been shown to successfully decode environmental sounds from functional magnetic resonance imaging (fMRI) data [16]; computationally demanding time-varying MPS features are decodable from spatially limited intracranial recordings [17]. The third set, *spectrogram*, provides a rich time-evolving frequency representation that was extracted for the same frequency bands as in FFT in 10 ms time windows, and could be linked with the high-dimensional MEG data in a computationally feasible design. In addition to these three models that include the fine spectral detail of the sound, we investigated decoding using the *amplitude envelope* alone without frequency information.

Time-sensitive decoding of the acoustic features was approached with a *convolution model* that convolves the MEG time series (amplitude of the evoked responses at each time point and sensor / cortical location) with a spatiotemporal response function learned from training data, such that the amplitude at each time-point *t* of the spectrogram (at 10 ms resolution) is reconstructed based on a corresponding time window in the MEG signals, separately for each frequency band *f.* The convolution is derived by integrating the time-locked MEG responses over a given lag window (*t − τ_2_*) to (*t − τ_1_*), where *−τ_1_*> 0, *−τ_2_ ≥* 0, and *t − τ_1_*> *t*, *t − τ_2_ ≥ t* (**Equation 1 in Methods; Fig. 1b**). Thus, the convolution model predicts spectrogram at time *t* using neural responses at time (*t − τ_2_*) and at successive 10 ms time points until time (*t − τ_1_*; see **Fig. 1b**, where *−τ_1_* = 180 and *−τ_2_*= 100). Thus, the lag values from stimulus to neural response are always positive (or zero), and the model incorporates the assumption that the neural activation follows (and never precedes) in time each time point of the stimulus that it is encoding. A short lag window entails that the activation of neuronal populations reflected in the MEG signal falls and rises closely following (i.e. time-locked to) the amplitude fluctuations within different frequency bands. The unknown weights for reconstructing the amplitude at each frequency band of the spectrogram were estimated by minimizing the regularized sum of squared error between the original spectrogram and the predicted sound spectrogram.

The time-averaged frequency / modulation content of the sounds, estimated with FFT / MPS respectively, was decoded using a *regression model*, in which each feature of the stimuli is predicted based on all time points (or a selected time window) of the MEG responses (**Equation 4 in Methods; Fig. 1b**) [36]. The unknown weights for reconstructing power at each frequency band / rate-scale-frequency point were learned in a similar fashion as for the convolution model; however, as these acoustic features (FFT/MPS) are non-time-varying, the whole MEG response or a selected time window in the response is used to predict them. Thus, both the regression and convolution models decode acoustic features from multivariate spatiotemporal neural patterns, but the regression model does not model the potential temporal tracking of stimulus features in the neural responses.

The FFT features predicted the MEG responses for spoken words reasonably well (average item-level decoding accuracy 65%, p < 10^−14^; **Fig. 1c**). Use of MPS resulted in similar performance (65%, p < 10^−11^; FFT vs. MPS, two-tailed Wilcoxon signed rank test, Z = 0.10, p = 0.94). In these regression models, MEG data at 0 – 1000 ms from stimulus onset was used to obtain an overview of the model performance. In contrast, the time-sensitive convolution model was remarkably successful at predicting spoken word spectrograms (83%, p < 10^−16^), with a significant improvement in comparison to the regression model that used time-averaged MPS (Z = 3.5, p = 0.000031; **Fig. 1c**); to obtain an overview of the convolution model’s performance, we used a lag window of 0 – 420 ms (delay from time point in the spectrogram to a range of time points in the MEG signal).

The success of the spectrogram convolution model in predicting spoken words was not merely due to increased information content (e.g. number of features) of the time-varying (spectrogram) compared to the time-averaged (FFT/ MPS) feature sets: When we used a regression model to predict the spectrogram frequencies of spoken words by including all time-points of the MEG response to predict each time point in the spectrogram, the convolution model continued to perform significantly better (spectrogram regression 73%, spectrogram convolution 83%; Z = 3.3, p = 0.00031; **Fig. 1c**). Thus, the salient improvement of predictive accuracy with the spectrogram convolution model suggests that the neuronal population activity in the auditory cortices closely follows the speech signal in time, to accurately encode the minute changes within spoken words.

We then investigated whether changes in the amplitude envelope of the spoken words, corresponding to the slow temporal modulations within the speech rhythm, are important for the remarkably successful decoding of spoken words, by separately decoding the amplitude envelope (spectrogram averaged across frequency) with the convolution model. The results reveal remarkably high classification performance for the spoken words (91%, significant difference from spectrogram decoding for speech, Z = 3.5, p < 0.001; **Fig. 1c**), further highlight the importance of the temporal aspects of the stimulus for speech decoding.

### Time-locked encoding of spoken words reflects acoustic-to-phoneme mapping

Leave-one-out-reconstruction of the spoken word waveforms with the spectrogram convolution model suggested preservation of acoustic properties characteristic of different phonemes (**Audio S1, Fig. S3**). To test this hypothesis, phonemic annotation of the speech sounds was aligned to the stimulus time-course [7, 21]. These *phoneme sequences* (**Fig. 1a**) were decoded successfully with the convolution model (73%; p < 10^−16^; **Fig. 1c**). Furthermore, a representation of the spoken words that combined both the speech spectrogram and the sequence of phonemes performed even above the spectrogram alone (84%; Z = 2.5, p = 0.01). Thus, information of the unfolding sequence of categorical phonemes is represented cortically along with the acoustic features of the spoken word.

To determine at what delay after each time point in a sound the brain responses reflect encoding of its acoustic and phoneme information, we investigated different lag windows between time points of the stimulus and the MEG response. For this analysis, we chose the spectrogram model, which allows for a reconstruction of the spoken words that retains the relevant sound features (in contrast to the amplitude envelope alone), and additionally analyzed whether phoneme information is represented at a similar lag. The lag window was advanced in non-overlapping 80-ms steps (20 – 100 ms, 100 – 180 ms, and so on until 340 – 420 ms). Both spectrogram and phoneme decoding with the convolution model performed best when MEG responses at a lagged windows of 100–180 ms after each time point in the sound were used (81% at 100–180 ms vs. 73% at 180–260 ms lag Z = 3.5, p = 0.000031; for phonemes 69% at 100–180 ms vs. 62% at 180–260 ms lag Z = 2.4, p = 0.016; **Fig. 2a**). For a sanity check, we also calculated the decoding accuracy for speech spectrogram using a counterintuitive lagged window of −80–0 ms, i.e. evaluating if “past” neural response could predict “future” spectrogram. With this lag, decoding performance was at chance level (mean decoding accuracy across 16 subjects 55%, SEM = 1.85). The cortical areas contributing to successful performance of the acoustic and phoneme models concentrated in and around the left and right auditory cortices (**Fig. 2b**).

**Figure 2.**
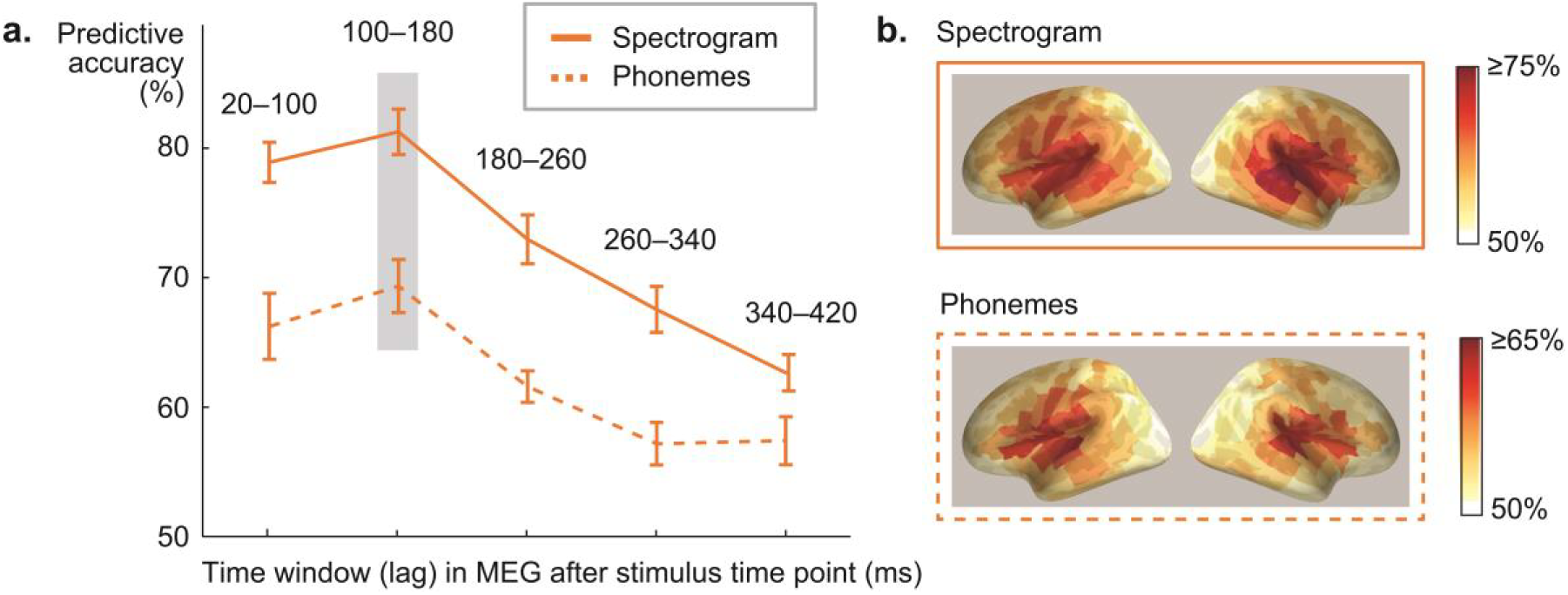
Influence of convolution lag and cortical sources for spectrogram and phoneme decoding. **a)** Spectrogram and phoneme decoding accuracy at different lags between a time point in the stimulus and a time window in the MEG response (average across all 16 participants +/− SEM). Note that the lag window does not correspond to timing relative to the stimulus onset in the MEG evoked response. The best predictive accuracy for decoding spectrogram and phoneme sequence of spoken words was reached with a lag of 100–180 ms (significant difference to 180–260 ms lag, p = 0.000031 for spectrogram, p = 0.016 for phonemes). **b)** Cortical sources contributing to decoding of acoustic and phoneme features in spoken words with the convolution model at 100–180 ms lag. Color scale denotes average decoding accuracy (> 50%) across all 16 participants.

To explore the contribution of semantics in the acoustic-phonetic decoding of spoken words, we investigated how these different models decoded pseudowords (included in the experimental sequence to increase participants’ attention; **Methods**), by training the models with all the meaningful spoken words, and using all possible pairs of the 8 pseudowords (28 combinations) for testing. Pseudoword decoding showed similar performance as decoding of the meaningful words (FFT 68%, MPS 66%, spectrogram convolution 86%, phoneme convolution 75%), with significant improvement for the spectrogram convolution compared to the MPS (Z = 3.4, p = 0.00015). The convolution model was able to predict the phoneme content of the pseudowords with a similar accuracy (75%) as for the real words. Thus, time-locked encoding does not seem to depend on lexical content, giving further support to the idea that it might reflect mapping of acoustic content to prelexical linguistic units.

### Decoding of environmental sound acoustics does not benefit from time-locked models

To investigate whether the convolution model performs better than time-averaged models for other sounds other than speech, we separately applied the different acoustic models to a variety of environmental sounds. Environmental sounds (44 items) from various semantic categories were chosen to ensure a large variability in both the acoustic and semantic features of the sounds (**Fig. S1**, **Table S1**). They conveyed the same meanings as the spoken words, thus the conceptual endpoint of neural processing is presumed to be the same. Also, their cortical responses were similar to spoken words (**Fig. S2**). However, the processing steps for accessing the meaning of environmental sounds are presumably different than for spoken words.

The FFT features predicted the MEG responses for environmental sounds fairly well (60%, p < 0.0001; **Fig. 3a**). Use of MPS resulted in better performance than FFT (70%, p < 10-15; FFT vs. MPS, Z = 3.5, p = 0.000031); such an improvement in decoding was not observed for spoken words (for spoken words FFT and MPS decoding were both at 65%). In contrast, the accuracy with spectrogram convolution (68%, p < 10^−4^) did not improve compared to the MPS result (Z = 1.1, p = 0.27), as was observed for words (spectrogram convolution decoding for words was at 83%, with significant difference between spoken words and environmental sounds: Z = 3.5, p < 0.0001). Using a regression model to predict the spectrogram frequencies of environmental sounds resulted in similar decoding performance (68%, Z = 0.18, p = 0.87), and was also fairly similar to spoken word decoding (spectrogram regression 73%; no significant difference between spoken words and environmental sounds: Z = 1.9, p = 0.058). No time-dependence was observed for environmental sounds using different lag windows (61% at 100–180 ms vs. 62% at 180–260 ms lag; Z = 0.052, p = 0.98; **Fig. 3b**). The amplitude envelope decoding showed some improvement for the environmental sounds compared to the spectrogram convolution decoding (Z = 2.3, p = 0.021), but was still low (72%) compared to the remarkably high decoding performance for speech amplitude envelope (91%; significant difference between spoken words and environmental sounds: Z = 3.5, p < 0.0001). Similarly to the convolution spectrogram decoding for speech, the cortical areas contributing to decoding environmental sound acoustics mainly concentrated in and around the left and right auditory cortices (**Fig. 3c**).

**Figure 3.**
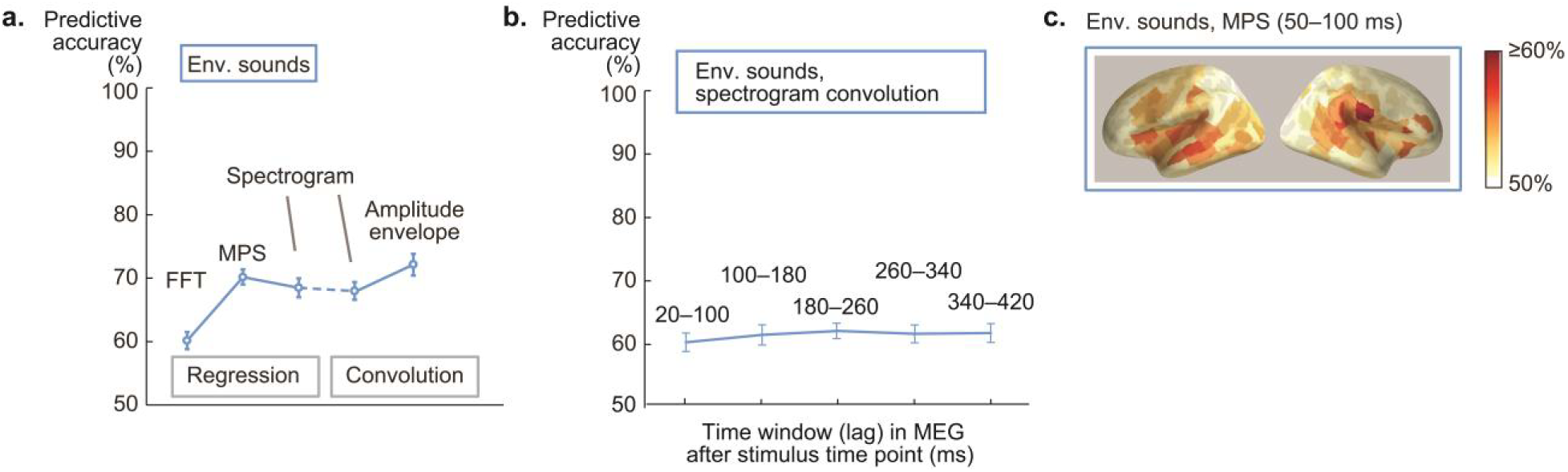
Comparison of different acoustic models for environmental sound decoding. **a)** Predictive accuracy for environmental sounds (blue) across all test sound pairs and 16 participants (mean ± SEM). The regression model was used for the decoding of non-time-varying features (FFT frequency bins / MPS rate-scale-frequency points / semantic features) and the convolution model was used for decoding of time-varying features (spectrogram time-frequency points / amplitude envelope); for a control analysis, the spectrogram was also decoded with the regression model. **b**) Investigation of convolution lag on spectrogram decoding accuracy for environmental sounds (average across all 16 participants +/− SEM) **c)** Cortical sources contributing to decoding of acoustic features in environmental sounds with the MPS model (at 50–100 ms after stimulus onset). Color scale denotes average decoding accuracy (> 50%) across all 16 participants.

### The observed enhancement for time-locked encoding of spoken words is not explained by possible confounds in the comparison of models

As the sounds were a selection of natural sounds, there were several differences between the spoken words vs. environmental sounds (**Fig. S4a, Fig. S5**). Thus, we sought to rule out potential confounds arising from these differences that might influence the observed striking improvement of decoding performance with the convolution model for speech but lack of similar improvement for environmental sounds.

We found that the benefit for spoken word but not environmental sound decoding was not due to larger dissimilarity among the spoken word stimuli with added temporal detail: While the spectrograms of spoken words do show larger dissimilarity than the MPSs (two-tailed t-test: t(945) = 139.5, p < 0.001), this is also true for the environmental sounds (t(945) = 14.8, p < 0.001; **Fig. S4b**). Notably, the dissimilarity among items, which typically leads to better decoding performance, is overall significantly greater for the environmental than speech stimuli, also with the spectrogram model (t(945) = 22.5, p < 0.001). This confirms that increased stimulus dissimilarity alone cannot explain the significantly better classification performance for spoken words than environmental sounds.

Furthermore, the benefits of the convolution model for modeling the cortical encoding of spoken words are not restricted to the leave-two-out classification task: Direct leave-one-out-reconstruction (correlation of the original and reconstructed features) demonstrated that spoken words were better reconstructed with the spectrogram convolution than MPS regression model (0.19 vs. 0.10, Z = 3.5, p < 0.001), whereas environmental sounds were better reconstructed with the MPS regression than spectrogram convolution (0.14 vs. 0.08, Z = 3.5, p = 0.001; **Fig. S4c**; see **Fig. S3** and **Audio 1** and **2** for examples of reconstructed sounds). The amplitude envelope of spoken words was also remarkably well reconstructed in comparison to the amplitude envelope of environmental sounds (reconstruction accuracy for spoken words 0.67 and for environmental sounds 0.32; Z = 3.5, p < 0.001; **Fig. S4c**). Moreover, correlating the pair-wise reconstructed spectrogram distances with the distances between original sound spectrograms shows that relatively more information is preserved in the reconstructions of spoken words compared to environmental sounds based on their respective cortical responses (Spearman correlation *r* = 0.41, p < 0.001 for spoken words, *r* = 0.22, p < 0.001 for environmental sounds; **Fig. S4d**).

The improved performance for spoken words was also unlikely to be solely due to different kinds of temporal properties of the words and other sounds. Specifically, speech has prominent slow (1– 7 Hz) temporal modulations (**Fig. 4a&b, Fig. S4a, Fig. S5**) that are important for its intelligibility [42] and have been suggested to be represented in the auditory cortices through a linear coding scheme [17]. Environmental sounds produced by the human vocal tract (laughter, crying etc.) are very similar to speech in terms of spectrotemporal characteristics and temporal modulation rates (with a small difference observed only for the highest rate (t(50) = 4.5, p < 0.001; Bonferroni corrected significance limit 0.0125; **Fig. 4c**). Spoken words showed a more variable spectral structure across time (larger values of SSI, Spectral Structure Index; see **Methods**) than environmental sounds (t(86) = 3.9, p < 0.001). However, the SSI did not differ between spoken words and human nonverbal sounds (t(50) = 0.45, p = 0.65), which were similar to each other also in their co-modulation properties (correlated temporal modulations between frequency channels; see **Fig. 4d**). Yet, we saw no improvement in decoding of this subset of environmental sounds with the convolution model (MPS regression 68% vs. spectrogram convolution 69%; Z = 0.31, p = 0.84). The results indicate that particularly spoken words, with their specific combination of spectrotemporal features, show remarkably improved decoding when neuronal population activity in the auditory cortices is modeled as tightly following the time-evolving acoustics.

**Figure 4.**
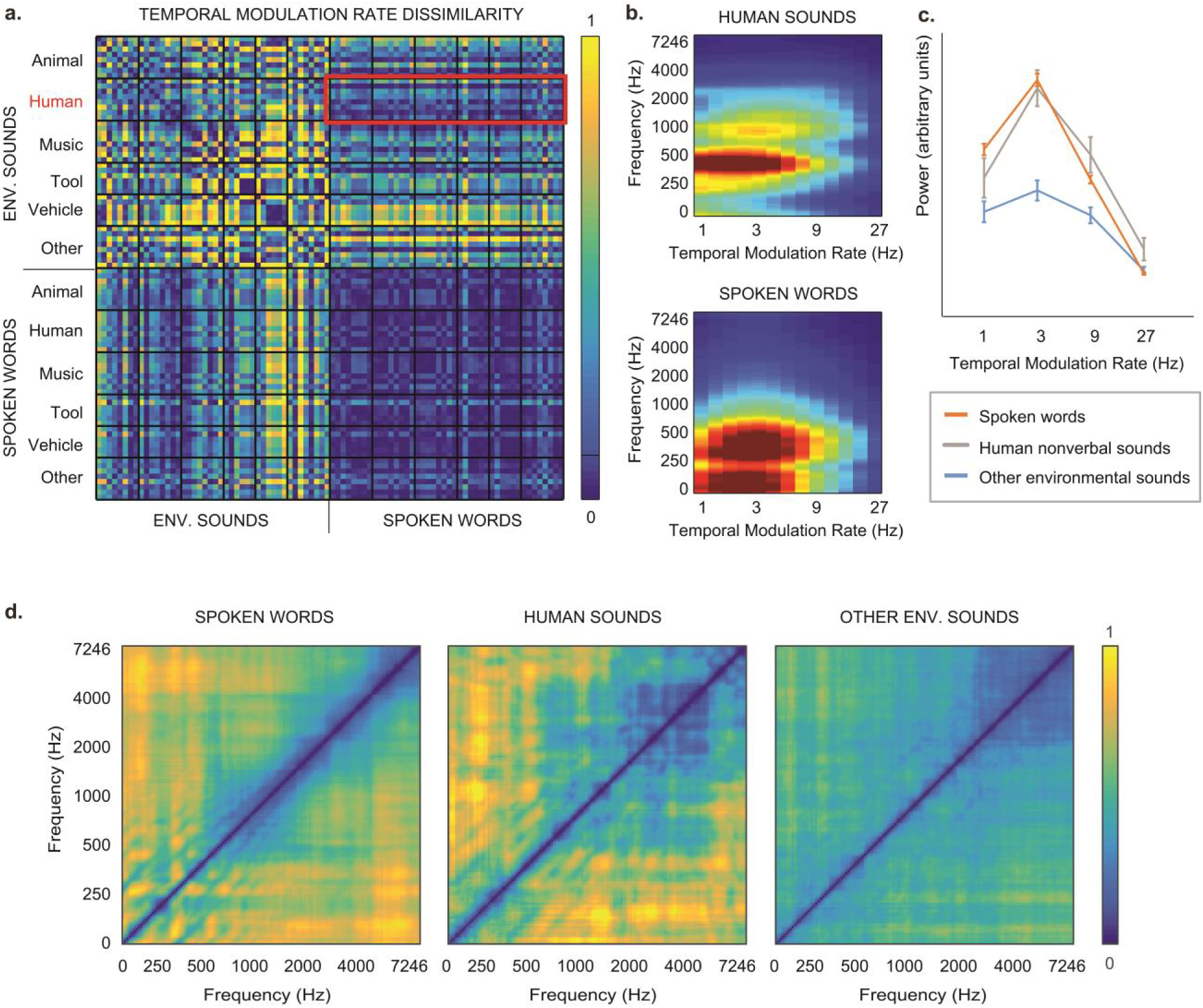
Comparison of human nonverbal sounds and spoken words. **a)** Dissimilarity (1 minus correlation) of the MPS temporal rates across the environmental sounds and spoken words. **b)** Average temporal modulation rate x frequency representations for human sounds (top) and spoken words (bottom). **c)** Power (mean ± SEM) at different temporal modulation rates for spoken words (red), human nonverbal sounds (grey) and other categories of environmental sounds (blue). **d)** Dissimilarity of different frequency bands of the sounds, indicating the degree of correlated temporal modulations (co-modulations), calculated separately for each item and averaged over spoken words (left), human nonverbal sounds (middle), and other environmental sounds (right).

### Semantic category is successfully predicted for both spoken words and environmental sounds, but with different timing

To ensure that the sounds were processed up to their meaning, the participants’ task was to identify immediate repetitions of the same meaning, i.e. two different exemplars of each sound, presented one after the other. The task was easier for the spoken words than for the environmental sounds (average percentage of hits for spoken words 94%, for environmental sounds 78%; two-tailed Wilcoxon signed rank test, Z = 3.6, p = 0.000031). The same set of semantic features (**Fig. 1a**, right) was used for modeling the environmental sounds and spoken words. The semantic features were obtained by combining two sets of norms, one acquired through a questionnaire and the other using word co-occurrences in a large-scale text corpus [14, 36]. The semantic feature representations captured meanings of individual items and formed salient clusters of the five semantic stimulus categories. We focused on predicting the semantic category (**Methods**). The regression model was successful in predicting the semantic category of both environmental sounds (81%; p < 10^−15^) and spoken words (58%; p = 0.0015), with significantly better accuracy for environmental sounds (Z = 3.5; p = 0.000031). Together, results of the behavioral task and the semantic decoding results indicate that attentional engagement was equal or even stronger for the environmental sounds than spoken words, and thus cannot explain the more successful acoustic decoding for spoken words than environmental sounds.

Semantic decoding reached significance for environmental sounds at 50–100 ms, and remained significant (p < 0.01) from 150 ms on until the end of the analysis window (**Fig. 5a**); we analyzed sensitivity of successive 50-ms time windows in the MEG responses. In contrast, MPS-based decoding of environmental sounds was significant at 50–150 and 200–300 ms after stimulus onset for environmental sounds and at 250–300 ms for spoken words. For spoken words, best performance for decoding spoken word semantics was reached late, and was significant only at 650–700 ms (**Fig. 5a**). Cortical sources contributing to semantic decoding are illustrated in **Fig. 5b** (see also **Fig. S6**).

**Figure 5.**
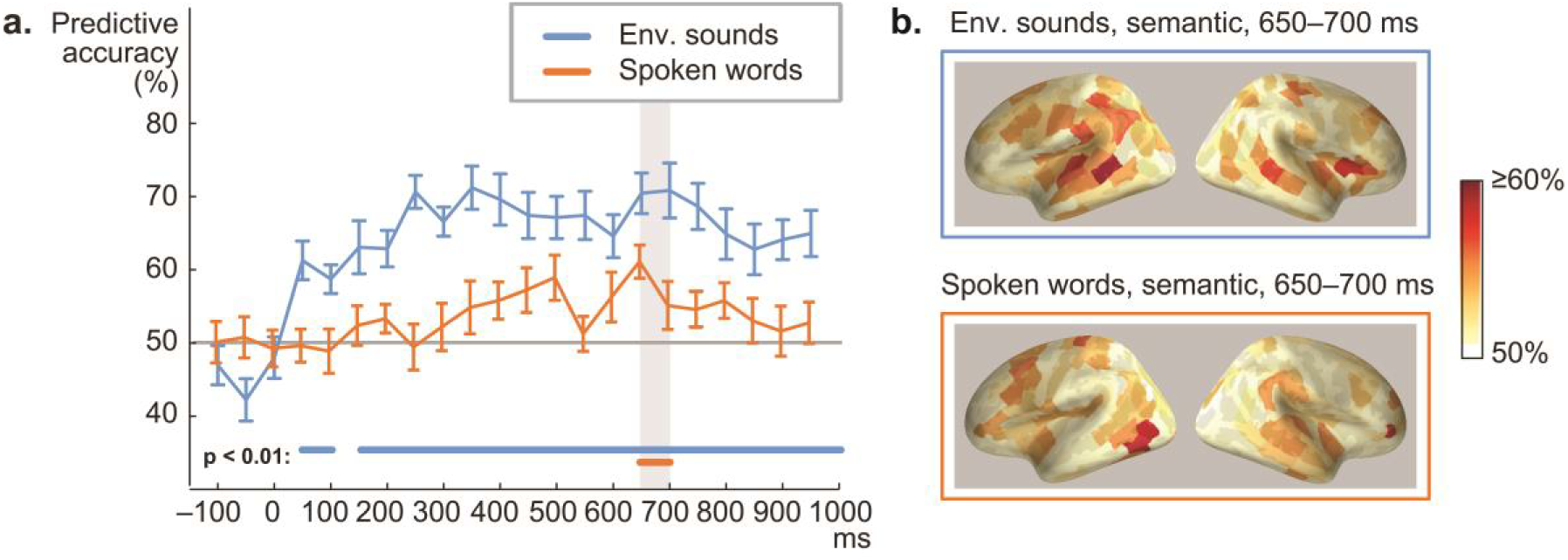
Comparison of semantic decoding for spoken words and environmental sounds. **a)** Time course of regression-model based predictive accuracy in semantic features of environmental sounds and spoken words (16 participants, mean ± SEM). The values on the x-axis represent the starting time of the 50-ms time windows in the MEG responses. Time 0 indicates the onset of the sound stimulus. Time windows with statistically significant decoding (p < 0.01) are indicated by thick horizontal lines above the x axis. Grey solid line denotes chance level performance (50%). The grey bars represent the time windows for source level decoding (section c). **b)** Source areas contributing to decoding of semantic features for environmental sounds and spoken words at 650 – 700 ms from stimulus onset.

## Discussion

### Cortical activation faithfully tracks the spectrotemporal detail of spoken words

The cortical processing of speech as compared to the processing of other sounds has remained a major open question in human neuroscience. Here, by combining time-sensitive brain imaging and advanced computational modeling, we discovered that the acoustic-phonetic content of natural, meaningful spoken words is encoded in a special manner, where the cortical evoked responses faithfully track in time the unfolding spectrotemporal structure as well as the amplitude envelope of the spoken words. This time-locked encoding was observed also for meaningless pseudowords. However, responses to environmental sounds, even human-made non-speech sounds with spectral and temporal modulations comparable to speech, did not show improved decoding with the dynamic time-locked mechanism, and were better reconstructed using the time-averaged spectral and temporal modulation content, suggesting that a time-averaged analysis is sufficient to reach their meanings.

These findings are highly relevant in view of the proposed role of envelope entrainment in speech processing. Phase-locking of oscillatory activation to the amplitude envelope of sounds has recently gained wide interest as a potential mechanism for cortical encoding of speech [25, 26]. However, it has remained controversial whether entrainment is directly related to speech parsing: It might reflect several different underlying processes [25], some of which are not specific to speech nor to the human auditory system [43]. Based on the current results, cortical auditory evoked activation tracks fine-scale spectral detail of single spoken words in a temporally accurate manner, relying heavily on both the spectrotemporal and overall amplitude changes within the sound. We propose that this time-locked mechanism may, at least partly, account for observations of oscillatory entrainment to continuous speech envelope [44]. Phase-locking of oscillatory activation to the stimulus may be driven by acoustic edges at syllable onsets [45] and can be modelled as a series of transient responses to changes in the stimulus [46]. Furthermore, the spectral content and temporal modulations within speech are non-independent, and envelope entrainment also depends on fine spectrotemporal detail [29]. Thus, phase-locked oscillations can be thought of as reflecting the superposition of transient evoked responses that track the fine-scale spectrotemporal evolution of the speech stimulus. Here, for evoked responses to isolated spoken words the best prediction was obtained for the amplitude envelope of the sound (reconstructed with 67% accuracy), indicating that the responses carry a lot of the characteristics of the envelope, possibly notably reflecting the sound onsets. Instead, for low-frequency cortical oscillatory activation to continuous speech, the best model appears not to be the amplitude envelope, but a combination of spectrotemporal features and phonetic-feature labels [21]. These acoustic features within speech are temporally coupled, and the grouping of features in the spectro-temporal fine structure is thought to be modulated by cues in the envelope [47].

Oscillatory entrainment relies on slow-rate temporal modulations of the speech envelope [48], which have been shown to be important for speech comprehension [42]. Notably, in our study, human vocalizations—the environmental sounds most similar to speech in that they contain prominent slow temporal modulations as well as similar co-modulations across frequencies—did not show any improvement in decoding with the time-sensitive spectrogram model. The dynamic mode of encoding uncovered here thus seems to take place for the particular combination of spectrotemporal features that are characteristic of speech. The high performance (83%) of the spectrogram convolution model, in which each time-frequency-point was predicted based on the MEG signal, showed that the fine spectral content of speech is tracked alongside the amplitude envelope at a high temporal resolution, and the spoken word spectrograms can be fairly accurately reconstructed based on this model. Our results, along with recent findings based on decoding sounds from fMRI responses [18], indicate that human auditory cortical processing might be optimally tuned for encoding the spectrotemporal structure of speech.

The current results corroborate previous work highlighting the importance of temporal modulations within speech [9], but they should not be interpreted to mean that auditory cortical activation would not be time-locked to non-speech sounds. Non-speech sounds also elicit prominent cortical auditory evoked responses time-locked to the acoustic changes within the sound, and phase-locking of cortical oscillations to the amplitude envelope is observed also for non-speech sounds [e.g. 32]. The current results further show that the amplitude envelope of environmental sounds is tracked in cortical evoked responses, as it can be reconstructed at 32% accuracy with the convolution model. Thus, a part of the time-locking phenomenon seems to be non-speech-specific. However, overall the best reconstruction of the environmental sounds (14%) was achieved by modeling their time-averaged modulation content (MPS), suggesting that the time-averaged spectral and temporal modulation content may be the information most accurately represented in cortical activation and sufficient for accessing meanings of nonspeech sounds. Indeed, the identification of common sounds in the environment may be based on the encoded summary statistics [16, 49]. Interestingly, the overall modulation content was a more accurate model of environmental sounds encoding than the frequency spectrum alone, whereas for speech there was no such improvement with modulation filters, consistent with a recent fMRI study [50].

For mapping speech acoustics to linguistic representations, accurate encoding of the temporal evolution within each frequency band is needed. An interesting parallel can be found from animal studies, where single-unit recordings in the auditory cortex suggest frequency-bin selective synchronization of neuronal population discharges to the temporal envelope of species-specific calls, which in many ways resemble human speech; thus, temporal envelope information within different frequency bands in behaviorally relevant vocalizations might be encoded cortically by coherent discharge patterns in distributed neuronal populations [51-53]. The current results suggest that, in humans, similar encoding mechanisms might have become especially important for speech, with extensive exposure to the language environment during development. Future studies should determine whether similar time-locking might be observed with tasks requiring attention to fine-grained temporal detail or specialization to categorical perception in other sounds through experience, e.g. for instrumental sounds in musicians.

### Accurate temporal encoding of the acoustic stream is necessary for linguistic parsing and semantic access of spoken words

Despite the different acoustic content of spoken words and corresponding environmental sounds, their processing converged at the endpoint; the same set of semantic features was successful in decoding both classes of sounds. Spoken words showed relatively low semantic decoding performance, in line with previous studies [22, 24]. One reason for the different semantic decoding performance of spoken words and environmental sounds may have been the fact that environmental sounds are inherently less familiar and their semantic access may thus require a more active effort than for spoken words, also suggested by the performance in the one-back task (detecting two subsequent sounds with the same meaning) which was more effortful, with more errors, for environmental sounds than spoken words. Nonetheless, our set of environmental sounds was tested for fast and consistent naming. In any case, the present finding of a striking differential effect of time information in the acoustic encoding of words, but no similar enhancement observed for non-speech sounds, is not likely due to increased attention to acoustic detail for the spoken words, as in fact more effort was needed to decipher the meanings of the environmental sounds. Semantic decoding of environmental sounds first reached significance at 50–100 ms, at the same time as acoustic decoding, whereas semantic decoding of spoken words reached significance clearly later, at 650–700 ms (this one significant time bin likely represents only a part of a wider time range). The current results are complementary to traditional experimental paradigms of semantic processing, where the violation of semantic expectation for spoken words and sounds shows in cortical responses at around 200–700 ms; these previous experiments have also indicated similar cortical effects of the semantic manipulations for spoken words and environmental sounds [54-56]. The current decoding results indicate, that for environmental sounds the acoustic features are informative of the physical sources (i.e. meaning) of the sound early on, whereas for spoken words intermediate phonological analysis steps have to be completed first. These include, according to the Cohort model, the comparison of the incoming input signal to the lexical representations stored in memory in a continuous manner, and selection of the correct lexical candidate [57]. For our stimulus words, which contained compound words, the uniqueness point and selection of one lexical candidate for semantic access was fairly late (on average 500 ms), which may have contributed to the late time window for the semantic decoding.

Previous studies have shown that the superior temporal areas are tuned to phoneme categories, which rely on integration of several spectral and temporal cues [7, 20]. In the current results, spoken word decoding improved compared to the spectrogram, when we included a combination of the spectrogram and the phoneme sequence of the word, with 100–180 ms the best lag for decoding both, suggesting that abstract phoneme information is represented in the cortical activation sequence simultaneously with the sound acoustics. The results echo those of a recent EEG study, where phoneme information and the spectrogram together were the best model for decoding continuous speech based on phase-locked cortical oscillatory activation, at a latency of 150 ms [21]. Several studies indicate that the optimal temporal integration window for parsing speech acoustics into linguistic units might lie within this range. The amplitude envelope of continuous speech is tracked by phase-locked cortical oscillations at latencies of about 100–180 ms [46, 58, 59], and this has also been shown to be approximately the time frame in which the categorical neural organization for phonemes transiently emerges [8]. Also, studies with isolated speech syllables have demonstrated that the evoked responses can be seen as a combination of transient ‘impulse responses’ to the onsets of constituent phonemes, with latencies of about 100–200 ms [60, 61]. However, the performance of the convolution model for spectrogram and phoneme decoding was fairly high also at 20-100 ms lag, suggesting that the acoustic-phonetic content of the words might in fact be tracked over multiple different integration windows. This may reflect the encoding of phonemes with different temporal characteristics [62], but may also be related to simultaneous phoneme and syllable level encoding, similarly to what has been observed in entrainment of nested cortical oscillations [25, 26].

The current results are compatible with the idea that abstract phoneme categories emerge directly from tuning to the complex spectrotemporal acoustic features characteristic of different phonemes [7, 50], and that the quasi-rhythmic changes in the amplitude envelope at syllable boundaries might be crucial in guiding this process. As noted above, this processing step where the continuous acoustic signal interfaces with phonological representations is only required for speech, and this is where time-locked encoding for speech seems to come into play. Decoding performance for the spectrogram and phonemes was similar for pseudowords and real words, indicating that this mode of encoding is related to the prelexical, low-level acoustic-to-phoneme conversion, and might not be directly affected by the lexical-semantic status of the speech utterance. However, this warrants further examination; a recent fMRI study showed that not only the spectral content and phonemic features, but also semantic features within narrative speech are represented in highly overlapping regions, early in the acoustic processing hierarchy [50], indicating top-down influences of semantic content on acoustic and phonological processing. In the current study, cortical sources contributing to acoustic and phoneme decoding were both concentrated around bilateral auditory cortices and did not show clear lateralization, consistent with previous studies [40, 50] and compatible with the view that acoustic-phonetic processing is bilaterally implemented [6].

### Future directions

The present discovery of a dynamic, time-locked mechanism for speech processing deepens the understanding and enables explicit modeling of the computations required for mapping between acoustic and linguistic representations. The current findings raise the question of what specific aspects within sounds are crucial for cueing the brain into using this special mode of encoding. Future work could investigate the contribution of different statistical properties within speech acoustics by using synthetized stimuli [9], the possible effect of experimental task to boost the use of time-locked or time-averaged mode in sound processing, and the role of top-down semantic contributions using real-life like auditory environments. The rapidly computable convolution model for high-dimensional MEG/EEG signals can be further developed for decoding of a time-varying modulation representation, which might even better model the time-locked cortical encoding of speech [17]. Finally, current findings offer a crucial link between investigations of cortical processing of continuous speech and isolated linguistic stimuli.

## Acknowledgments

We thank Mia Liljeström and Pekka Laitio for providing customized code for the source parcellation, Ossi Lehtonen for assisting with the source modeling, Jan Kujala with assistance in sound feature analysis, Tiina Lindh-Knuutila for assistance with the corpus vectors, and Sasa Kivisaari for comments on the manuscript. This work was supported by the Academy of Finland (grants 255349, 256887, 292552 and 315553 to RS, 277655 to HR), the Finnish Cultural Foundation (to HR), the Sigrid Jusélius Foundation (to RS), Maastricht University (to EF), the Dutch Province of Limburg (to EF), and the Netherlands Organization for Scientific Research (NWO grant 453-12-002 to EF), the Doctoral Program Brain and Mind (to AN), the Foundation for Aalto University Science and Technology (to AN) and Emil Aaltonen Foundation (to AN).

## Author contributions

A.N., A.F., J.S., H.R. & R.S. designed the experiment. A.N. & J.S. collected the data. A.N., A.F. & J.S. analyzed the data. A.N., A.F., J.S., H.R., E.F. & R.S. wrote the paper. E.F. gave technical and conceptual advice. R.S. supervised the project.

## Methods

### Data availability and contact for resource sharing

Ethical restrictions imposed by the Aalto University Research Ethics Committee prevent the authors from making brain imaging data publicly available without restrictions, as this data cannot be fully anonymized. Making the data freely available under the Creative Common licence would not enable us to restrict its use to scientific purposes. However, the relevant data used in this study are available from the authors upon reasonable request and with permission of the Aalto University Research Ethics Committee, for researchers aiming to reproduce the results. Further information and requests for resources should be directed to and will be fulfilled by the Lead Contact, Anni Nora.

### Participants

The human participants were 16 (8 male, 8 female) right-handed Finnish-speaking volunteers aged 19–35 years (mean 24 years). Exclusion criteria included hearing disorders and developmental or acquired neurological or language disorders. Participants signed a written consent form before the measurement, in agreement with the prior approval of the Aalto University Research Ethics Committee.

### Stimuli and experimental procedure

High-quality environmental sounds were chosen from internet sound libraries and, using the Adobe Audition program, modified to readily identifiable mono sounds of approximately 1 s duration, with sampling frequency 44.1 kHz and bit rate of 16 bits. With careful selection of sounds from several categories, we sought to include as much acoustic variability as possible to extensively span the low-level acoustic feature space. The experimental stimuli were selected from these sounds based on a behavioral test with 12 participants: Participants wrote down the name of each sound immediately after recognizing it. Environmental sounds with at least 80% consistent naming and response time less than 3 s were chosen as stimuli. The final 44 environmental sound stimuli belong to 6 categories: *animals*, *human sounds*, *tools*, *vehicles*, *musical instruments* and *others* (6–8 items per category; see **Fig. S1**, **Table S1**). The *others* category included items that did not belong to any of the other above-mentioned categories. The same environmental sounds were used as stimuli across all participants.

The spoken word stimuli (44 in total) were the noun labels of the final selection of environmental sounds. To increase acoustic variability, each word within a category was spoken by a different speaker (8 speakers in total; 4 female, 4 male; two children/adolescents). Also, each speaker spoke one word in each semantic category, to enable eliminating the influence of speaker-specific acoustic features from category-level decoding. The speaker set was rotated across participants. Stimulus duration was on average 810 ms (s.d. 180 ms) for the spoken words and 920 ms (s.d. 230 ms) for the environmental sounds. The uniqueness points (i.e. estimated time of lexical selection) was on average about 500 ms (range 300–890 ms). Due to the transparent nature of Finnish the uniqueness point of a spoken word (point of divergence from all other words with a different word stem) corresponds to its orthographic uniqueness point, and thus can be calculated in a straight-forward fashion. Here it was calculated based the same Finnish Internet-derived text corpus, that was used to create the semantic features [63]. All sounds were filtered with an 8-kHz linear low-pass FFT filter (Blackman-Harris) and re-sampled at 16 kHz. Mean amplitudes of the stimuli were normalized such that the root-mean-square power of each stimulus was the same. Before the MEG measurement, the individual hearing threshold was determined for each participant, and the stimuli were delivered through plastic tubes and earpieces at 75 dB (sensation level).

To reach a high enough signal-to-noise ratio (SNR) per stimulus item for the machine learning approach, each stimulus was presented 20 times in a pseudorandom manner, such that two words spoken by the same speaker, or a spoken word and an environmental sound referring to the same meaning were not presented in a row. The participants’ task was to listen carefully to each sound, think about the concept it refers to, and respond with a finger lift when two sounds with the same meaning were presented one after another (one-back task, 4% of trials). Response hand was alternated between participant pairs. In the task trials, another exemplar for each word or environmental sound was presented as target stimulus (same word spoken by a different speaker or same environmental sound with a different acoustic form, e.g. a different kind of dog bark; the two stimulus types were not mixed in this task, e.g. the word cat was never followed by a cat sound).

Additional filler items, 8 meaningful environmental sounds and 9 spoken words, were included to make the sequence more variable. In addition, 8 meaningless environmental sounds and 8 pseudowords were presented 20 times each to increase participants’ attention. The meaningless environmental sounds were unidentifiable sounds from the abovementioned categories, which were further processed by reversing them in the time domain or scrambling them in 50 – 100 ms segments. The resulting sounds shared properties with the meaningful sounds from the same categories, but they were not identifiable in the behavioral pre-test. The pseudowords were minimal pairs to real Finnish words, following Finnish phonotactic rules. The task trials, filler sounds and meaningless sounds were excluded from the main analysis. However, we separately investigated pseudoword decoding with the different models to explore the possible contribution of lexical and semantic aspects in spoken word decoding.

### Acoustic, semantic and phoneme features of the sounds

#### Acoustic features

We used three different sets of acoustic features that included spectral detail of the sounds: Frequency spectrum, Spectrogram (time-varying frequency spectrum) and a time-averaged Modulation Power Spectrum (MPS). The frequency spectrum is a non-time-varying representation of the stimulus power per frequency. Fast Fourier transform (FFT) was calculated over the entire sound time window by using a filter bank of 128 frequency bands, with central frequencies of the bands ranging from 180 to 7246 Hz, uniformly distributed along a logarithmic frequency axis.

For a time-evolving representation of the acoustic features, a spectrogram was generated from sounds using an auditory filter bank with 128 overlapping band-pass filter channels, with their central frequencies corresponding to the FFT frequency bands [following 17], using the NSL-toolbox (http://www.isr.umd.edu/Labs/NSL/Downloads.html). This filter bank mimics the representation of sound in the human cochlea [15]. The time resolution and other parameters were optimized for retaining the characteristic temporal modulations of the sounds, while enabling extraction of the relevant spectral structure of the sounds. The sounds were divided into frames of 10 ms and integrated over 16-ms time windows.

Next, we calculated the modulation power spectra (MPSs) based on the spectrograms using the NSL-toolbox [15]. MPSs were calculated with modulation-selective filters spanning wide to narrow spectral scales (0.5, 1, 2 and 4 cycles/octave) and slow to fast temporal modulation rates (1, 3, 9 and 27 Hz) within the sounds (see **Fig. 4, Fig. S4**). These modulation selective filters mimic the spectrotemporal receptive fields of the primary auditory cortex [15], and have previously been shown to be successful in decoding and reconstructing natural sounds based on intracranial recordings [17] and fMRI data [16, 18]. The four temporal modulation rates and four spectral scales selected for the present study have been shown to capture the essential features of a broad range of natural sounds [16]. We chose a three-dimensional frequency-specific MPS where the time dimension of the MPS was averaged out, and upward and downward going modulation rates were averaged together, in line with a recent comparative evaluation of different MPS representations using fMRI [16]. The use of non-time-varying (time-averaged) MPS features offered critical savings in computational effort and also enabled direct comparison with the decoding of non-time-varying semantic features.

Finally, we investigated a representation of the sounds that contains temporal modulation, but lackes fine spectral structure, by using the amplitude envelope of the sounds. The amplitude envelope was estimated through averaging the sound spectrogram across the 128 frequency bins in 10 ms windows, resulting in one feature vector.

The number of acoustic features were 128 in FFT (128 frequency bands), on average 11100 in spectrogram (128 frequency bands × 87 time windows of 10 ms), 2048 in MPS (128 frequency bands × 4 spectral scales × 4 temporal modulation rates) and on average 87 in spectrogram (1 amplitude envelope × 87 time windows of 10 ms); for spectrogram and amplitude envelope, the length of the feature vectors varied with the sound length. To control for possible confounding effects of the varying lengths of feature vectors on the performance of the spectrogram convolution model, the spectrogram vectors for the two held-out test sounds were equalized to the length of the shorter one.

#### Phoneme sequence

Phoneme features of the words were obtained through their phonemic annotation, time-aligned to the stimulus wavefile [7, 21]. Each phoneme was set as 1 in those 10 ms time windows when it was present and otherwise as 0. There was no overlap, i.e. only one phoneme was marked “active” in each time window. Phonemes with 10 or more instances in the stimulus set were included, resulting in a set of 15 phonemes, each occurring 10–40 times. This representation had on average 1300 features.

#### Combined spectrogram and phoneme features

For investigating whether the spectrogram and phoneme decoding results were based on fully overlapping information, we combined these into a single feature representation of the spoken words, with on average 11100 spectrogram features plus 1300 phoneme features; the number of features varied with sound length.

#### Semantic features

Semantic features were obtained by concatenating two sets of semantic norms, corpus statistics and question norms. For extracting the corpus statistics, the frequencies of co-occurrences of words in the immediate neighborhood (5 words before and 5 words after) of each lemmatized stimulus word (compound markers added, some words changed from plural to singular) were calculated from a 1.5 billion-token Finnish Internet-derived text corpus [63] using a continuous skip-gram Word2vec-algorithm using default parameters [64]. The dimensionality of the trained vectors was 300, i.e. a vector with 300 values was used to describe each item.

Question norms for the stimulus words were collected with a web-based survey. The questions in the survey were partly based on a previous study [36] but modified to better represent the present selection of stimulus categories. Fifty-nine university students (32 female, 27 male, mean age 26 years; none of them participated in the present MEG study) answered 99 questions, presented in random order, about the semantic properties of each item on a scale from 0 to 5 (from definitely not to definitely yes); values for each question were averaged across the participants. Each item was thus described with a vector with 99 values.

#### Similarity measures

To investigate how the different acoustic models (especially the MPS and spectrogram models) differ in the dissimilarity among the speech vs. non-speech sounds, we have calculated the pairwise distances (1 minus sample correlation) for all spoken words and environmental sounds (**Fig. S4b,d**). We chose correlation as our similarity measure, as correlation between the original and reconstructed model of the sound was also used in the evaluation of the classification and reconstruction performance. Thus, this measure serves as a baseline measure of discriminability of the items with the different models, and for the spectrogram model it describes the cumulative dissimilarity of all time points at each frequency band of the spectrogram. To calculate the distances (1 minus correlation), the different frequency bands were concatenated.

#### Sound complexity measures

For estimating how the spectral distribution fluctuates across time in environmental sounds and spoken words we calculated, first, the variance of the spectral structure (spectral flatness measure SFM, an estimate of the number of distinct peaks in the frequency spectrum [65]) across each 10-ms segment within the spectrogram and, then, the spectral structure index (SSI, a measure of spectral variability defined as the SFM across time and suggested as a measure of sound complexity [65]). Larger values of SSI denote more variable spectral structure across time.

### MEG recording

Magnetic fields associated with neural current flow were recorded with a 306-channel whole-head neuromagnetometer (Elekta Oy, Helsinki) in the Aalto NeuroImaging MEG Core. The sensor array consists of 102 triple sensor elements, each with one magnetometer and two planar gradiometers. Planar gradiometers give maximum signal above current source, whilst magnetometers are sensitive to far away sources as well. The MEG signals were band-pass filtered between 0.03 and 330 Hz and sampled at 1000 Hz. Eye movements and blink artifacts were monitored by two diagonally placed electro-oculograms (EOGs). The position of the participant’s head within the MEG helmet was defined using five head position indicator coils. The locations of these coils, attached to the participant’s scalp, were determined with respect to three anatomical landmarks (nasion and two preauricular reference points) with a 3D digitizer, and with respect to the sensor array by briefly feeding current to the coils during the measurement. Head movements were monitored continuously [66]. The MEG measurement lasted approximately 40 minutes, with breaks every 5 minutes.

### Anatomical MRI acquisition

Anatomical magnetic resonance images (MRIs) were obtained with a 3T MRI scanner (Magnetom Skyra, Siemens) for all 16 participants. The scan included a 3-plane localizer and a T1-weighted anatomical image. To allow attribution of MEG activation patterns to cortical loci, the MEG data were co-registered in the same coordinate system with the individual MR images.

### MEG preprocessing and source modeling

Spatio-temporal signal space separation [tSSS; 67] and movement compensation algorithms [66] were applied offline to the raw data using Max-Filter™ software (Elekta Neuromag), to remove the effects of external interference and to compensate for head movements during the measurement. To obtain an estimate of the artifact signals caused by blinks or saccades, the MEG signals were averaged with respect to transient maxima in the EOG signal, principal component analysis (PCA) was performed on this average, and the corresponding magnetic field component was removed from the raw data [68].

The MEG data analysis focused on planar gradiometer channels. Single trials were averaged from 300 ms before to 2000 ms after the stimulus onset, rejecting trials contaminated by any remaining artifacts (signal strength exceeding 3000 fT/cm). On average 19.7 ± 0.6 (mean ± s.d.) artifact-free epochs (trials) per participant were gathered per item (maximum = 20). The averaged MEG responses were baseline-corrected to the 300-ms interval immediately preceding the stimulus onset. The averaged data was low-pass filtered at 40 Hz.

An estimate of the underlying cortical sources was obtained with minimum norm estimates [MNE; 69] using MNE Suite software package [70]. For MNE analysis, the cortical surface of each participant was reconstructed from the corresponding magnetic resonance (MR) images with Freesurfer software [71, 72]. Each hemisphere was covered with ∼5000 potential source locations. Currents oriented normal to the cortical surface were favored by weighting the transverse currents by a factor of 0.2, and depth-weighting was used to reduce the bias towards superficial sources [73]. Noise-normalized MNEs (dynamical Statistical Parametric Maps, dSPMs) were calculated over the whole cortical area to estimate the SNR in each potential source location [74]. Noise covariance matrix was estimated from the 300-ms prestimulus baseline periods across all trials. The source space (∼10000 vertices) was divided into 229 parcels of approximately equal size (101 in the left and 118 in the right hemisphere), using the Destrieux Atlas as a starting point [75]. Medial and orbitofrontal areas were excluded, as MEG does not reliably measure activation in these areas. Before applying the parcellation, the cortical surface of each participant was morphed onto Freesurfer’s average cortical surface template (fsaverage).

### Machine learning models

Before model fitting, MEG sensor level responses were downsampled to 100 Hz (i.e. sampled at 10 ms resolution). All decoding models were trained and tested separately for the spoken words and separately for environmental sounds, and for the sensor- and source-level data. The model training and testing was done separately within each individual participant. In the sensor space, the analysis was restricted to 28 planar gradiometer pairs over the bilateral auditory cortices for the acoustic and phoneme decoding, whereas signals from all 204 planar gradiometers were used for semantic decoding. For decoding of time-varying features, i.e. the spectrogram frequencies, amplitude envelope and phoneme sequences, we used a convolution model. For decoding of non-time-varying features, i.e. semantic features, frequency spectrum (FFT) and time-averaged MPS, we used a regression model.

Prior to performing machine learning analysis, the stimulus features and MEG signals were *standardized* across all stimuli, such that the mean value was set to zero and standard deviation to unity: For the acoustic features, the FFT and spectrogram were normalized within each frequency band, and the MPS within each rate, scale and frequency band. Each semantic feature was similarly standardized across all stimuli. The MEG signal power per sensor or source location was normalized within each 10 ms time window. When applied to both stimulus features and corresponding MEG responses, this procedure ensures that the absolute power per frequency band is not crucial in model estimation. Instead, the unknown quantities (weights in regression and spatio-temporal response functions in convolution) are estimated such that the variation in the MEG signal power consistently correlates with variation in each stimulus feature, across stimuli.

Data standardization is a normal practice in statistical analysis and ensures that individual data features are weighted equally in the analysis [76].

*Convolution model* [17, 33, 34] searches for a suitable mapping (stimulus response function) between the time-varying neural response and the time-varying stimulus features. Here we use a novel scalable formulation of the model that allows decoding of time-varying properties of the sounds (*s*) by convolving the time-varying MEG responses (*r*) at brain location *x* with an unknown spatiotemporal response function *g* [35]. The model predicts stimulus features (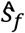, where *f* is a given stimulus feature, here the amplitude at one spectral frequency band or the value (0/1) of one phoneme) at time *t*, based on the MEG activation (*r*) integrated from time (*t − τ_2_*) to (*t − τ_1_*); *−τ_1_* > 0, *−τ_2_ ≥* 0 (see **Fig. 1b**, where *−τ_1_*= 180 and *−τ_2_*= 100). The procedure is repeated independently for all frequency bands of the stimulus, resulting in prediction of the time series of amplitude changes of sound spectrogram, amplitude envelope or the phoneme sequence,

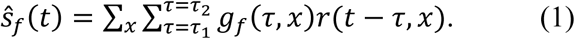

To estimate the unknown spatiotemporal response functions *g_f_* we use the dual representation of **Equation 1**, and impose an L2 prior on the unknown reconstruction weights, *g_f_*(*τ*, *x*) [35] as follows. Using matrix notation, **Equation 1** can be written as, *S_f_* = *RG_f_*, where we define *S_f_ ∈ ℝ*^(*NT*)×*1*^, *G_f_ ∈ ℝ*^(*τx*)×*1*^, and the response matrix *R ∈ ℝ*^(*NT*)×(*rx*)^, such that each row *τ_n_*(*t*) in *R* represents MEG response to a spoken word or environment sound *n* across all sensors *x* and all time points sampled from (*t − τ_2_*) to (*t − τ_1_*). The unknown *G_f_* are estimated by minimizing the regularized sum of squared error between the original s*_f_* and the predicted sound features 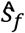,

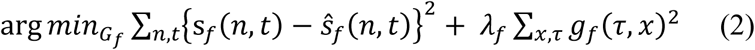

Minimizing this loss function leads to the maximum-a-posteriori (MAP) estimate for *G_f_*,

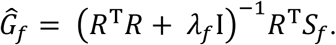

This classical MAP estimate is not ideal for MEG studies where the number of conditions N are typically small compared to the dimensionality of neural responses since *R^T^R ∈ ℝ*^(^*^τx^*^)×(^*^τx^*^)^ [34, 35]. Therefore, similar to kernel ridge regression [77] (also known as dual representation of ridge regression) we obtain the MAP estimate of the convolution model using its dual representation where the inner product *R^T^R* is replaced with corresponding Gram matrix *RR^T^*,

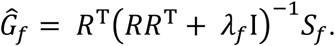

To estimate the regularization parameter *Ji_f_* we use a grid of pre-defined values for the hyper-parameter and choose the optimum value that minimizes leave-one-out error within training data [77]. Given the lag parameters *τ_1_* and *τ_2_* the learned MAP estimates *g_f_*(*r*, *x*) are used to predict the acoustic and phonetic features of a new, previously unencountered, test sound as follows:

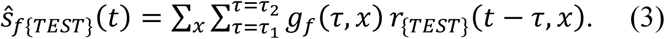

The convolution model thus predicts the spectrogram at time *t* using neural responses at time (*t − τ_2_*), …, (*t − τ_1_*). To obtain an overview of the model’s performance, we used a lag window of 0 – 420 ms (delay from time point in the spectrogram to a range of time points in the MEG signal, i.e. from *−τ_2_* to *−τ_1_*, where *τ_1_*= *−*420 and *τ_2_*= 0, an estimate based on our recent evaluation [35] (**Fig. 1c**). Next, we advanced the lag window in non-overlapping 80 ms steps (20 – 100 ms, 100 – 180 ms, …, 340 – 420 ms) to investigate the dependence of the decoding accuracy on the lag timing (**Fig. 2a**). A relatively large step size (80 ms) was chosen due to the high computational load of the analysis and also motivated by previous research showing oscillatory entrainment at latencies of about 100–180 ms [46, 58, 59].

In the *regression model*, the dependent (predicted) variables were the semantic or non-time-varying acoustic features, while the independent variables were the MEG responses *r(t,x)* at brain location *x* and time *t* from stimulus onset. Unknown weights *w_f_*(*t*, *x*) and the L2 regularization parameters were learned in a similar fashion as for the convolution model using dual representation of the regression model [36]. Using the same notation as in **Equation 1**, the reconstruction (predicted value 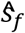) for one semantic or non-time-varying acoustic feature *f* can be written as:

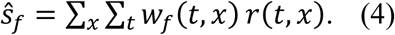

We first utilized MEG data at 0 – 1000 ms from stimulus onset to obtain an overview of model performance (**Fig. 1c**) and then analyzed sensitivity of successive 50-ms time windows (0–50 ms, 50–100 ms, etc.) in the MEG responses to see how the decoding of semantic/acoustic features varied with time (**Fig. 5a**). This resolution was considered sufficient to cover the early transient response components which were expected to be especially important for encoding the acoustic features [5]. In the source space, the models were learned, for each participant, separately for the vertices within each cortical parcel, and visualized by averaging the decoding performance within each cortical parcel across participants. The cortical sources contributing to acoustic and semantic decoding were visualized using the model and time window that performed best for each class of sounds (spectrogram-based convolution with 100–180 ms lag for acoustic and phoneme decoding of spoken words, **Fig. 2b**; MPS-based regression at 50–100 ms for acoustic decoding of environmental sounds, **Fig. 3c**; regression at 650–700 ms for semantic decoding, as this time window had significant decoding performance for both classes of sounds, **Fig. 5b**). We visualized the sensitivity of cortical areas in decoding stimulus features by averaging the decoding performance within each cortical parcel, across all participants. For investigating overlap of sources between acoustic and semantic decoding, we focused on the 20 highest ranking areas in terms of mean decoding performance (**Fig. S6**).

### Performance evaluation

We adopted a two-stage evaluation scheme [36, 37]. In the first stage, the convolution or the regression model was trained to learn the unknown weights using all but two held-out items/sounds. In the second stage the learned weights were used to decode stimulus features (acoustic, phonemic or semantic features) for the two held-out test items. The decoding was considered correct if the combined similarity between the true features of sounds (*s*1 and *s*2) and the decoded features of the sound (*p*1 and *p*2) was greater than the reverse labeling, i.e.

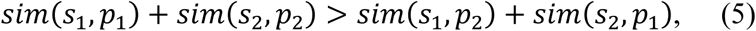

where *sim*(*s*, *p*) is the Pearson correlation between original *s* and decoded features *p*. The evaluation scheme of **Equation 5** was repeated for all possible combinations in a leave-two-out cross-validation approach as follows: For *item-level decoding* (for all acoustic and phoneme features), the 44 items (spoken words or environmental sounds) were divided into 42 training and 2 test sounds in all possible pair-wise combinations, leading to a total of 946 pair-wise tests. For *category-level decoding* (for semantic features), the two held-out test sounds were always chosen from two different semantic categories, resulting in 66 possible pair-wise tests. The evaluation was done separately for spoken words and environmental sounds. For spoken words, these test sounds were always chosen from the same speaker (i.e. *within-speaker prediction*); thus this category-level prediction is not influenced by speaker-related acoustic differences. The reported participant-level decoding accuracy is an average over all pair-wise combinations of the held-out pairs.

### Statistical significance

Statistical significance was established by repeating each analysis with permuted data, and empirical p-values were obtained for single participants. In each permutation run, data from one participant was chosen at random and the item labels for the averaged evoked responses were randomly permuted across the different sounds (within spoken words or environmental sounds). This procedure was repeated 200 times for each convolution model and 1000 times for each regression model. For each permutation, the models were evaluated using all possible pairwise tests in a leave-two-out cross-validation scheme. Empirical p-values were computed by calculating the number of times the permutation result was better than the observed decoding accuracy for each of the 16 participants. The p-values across all 16 participants were combined using Fisher’s method and corrected for multiple comparisons (over time in the reported time courses) using false discovery rate (FDR) at an alpha level of 0.01. Two-tailed Wilcoxon singed rank test was used for comparing the predictive performance between different models and sound classes across all 16 participants.

## Supplemental materials

**Figure S1, related to Fig. 1**

Examples of spoken words (left) and environmental sounds (right), one from each semantic category. For each sound, the signal waveform is depicted on top, with the three models of the sound below: frequency spectrum (FFT), spectrogram, and MPS (temporal rates and spectral scales shown here separately). Note the considerable variability in the spectral content and temporal evolution of the different sounds, especially for the environmental sounds.

**Table S1, related to Fig. 1**

Complete list of stimulus items.

**Figure S2, related to Fig. 1**

**a)** Sensor-level responses to spoken words (left, orange) and environmental sounds (right, blue), in one MEG sensor above the left and right temporal lobes, for one participant. The signals were averaged across 20 presentations of the same sound, from 300 ms before to 2000 ms after stimulus onset, and here also averaged over the different items within each semantic category (for visualization only). Both spoken words and environmental sounds elicited a typical time-sequence of activation, with a transient response at about 100 ms that was followed by a more sustained response from about 250 ms onwards and return to baseline after 1000 ms. **b)** Grand average (n=16) dynamical Statistical Parametric Maps (dSPMs) to spoken words (left) and environmental sounds (right) demonstrate that, for both types of stimuli, activation originated mainly in the bilateral temporal regions in the vicinity of the primary auditory cortex, with additional activation in inferior frontal areas, and the left hemisphere was highlighted particularly for spoken words in the later time window.

**Figure S3, related to Fig. 1**

Original and reconstructed spectrograms of selected spoken words (top) and environmental sounds (bottom) based on the convolution model. The corresponding sound files are in **Audio S1 and S2**.

**Audio S1, related to Fig. 1**

Sample reconstructions of a selection of spoken words, based on the convolution model and spectrogram. The original stimulus is presented first, followed by the reconstructed sound. For corresponding spectrograms see **Fig. S3.**

**Audio S2, related to Fig. 3**

Sample reconstructions of a selection of environmental sounds, based on the convolution model and spectrogram. The original stimulus is presented first, followed by the reconstructed sound. For corresponding spectrograms see **Fig. S3**.

**Figure S4, related to Fig. 3**

**a)** The median (with 25% and 75% interquartile ranges) power spectra (left), as well as spectral scales (middle) and temporal rates (right) pooled across the 128 frequency bins, for speech and environmental stimuli, illustrating the differences between spoken words and environmental sounds. Also, the environmental sounds have more variability in their spectral scales and temporal rates, whereas spoken words display prominent slow temporal modulations. **b)** Pair-wise distances (1 minus correlation) of the original stimulus MPS and spectrogram (mean ± SD). **c)** Leave-one-out reconstruction fidelity, i.e. correlations of reconstructed and original MPSs / spectrograms / amplitude envelopes for spoken words and environmental sounds (mean ± SEM). **d)** Scatterplot of the pair-wise distances (1 minus correlation) among original (x-axis) and pair-wise distances among reconstructed (y-axis) spectrograms.

**Figure S5, related to Fig. 3**

The median (with 25% and 75% interquartile ranges) power spectra (left), as well as spectral scales (middle) and temporal rates (right) pooled across the 128 frequency bins, for speech and different categories of environmental stimuli. The human nonspeech sounds are most similar to speech in the spectral scales and temporal rates.

**Figure S6, related to Fig. 5**

Illustration of the cortical parcellation template and the top 20 ranking cortical areas (parcels) for the decoding of acoustic (orange) and semantic (red) features for spoken words (left), and acoustic (light blue) and semantic (dark blue) features for environmental sounds (right). The parcel names are listed in the table below. Sources for acoustic decoding of spoken words are based on the convolution model and spectrogram, with the best lag (100–180 ms). Sources for acoustic decoding of environmental sounds are based on the regression model and MPS, at 50–100 ms. For semantic decoding, the regression model at 650–700 ms was used for both classes of sounds. L = left, R = right.

