## Supplementary figures and images for "Dynamic time-locking mechanism in the cortical representation of spoken words"

### Supplemental Figure 1

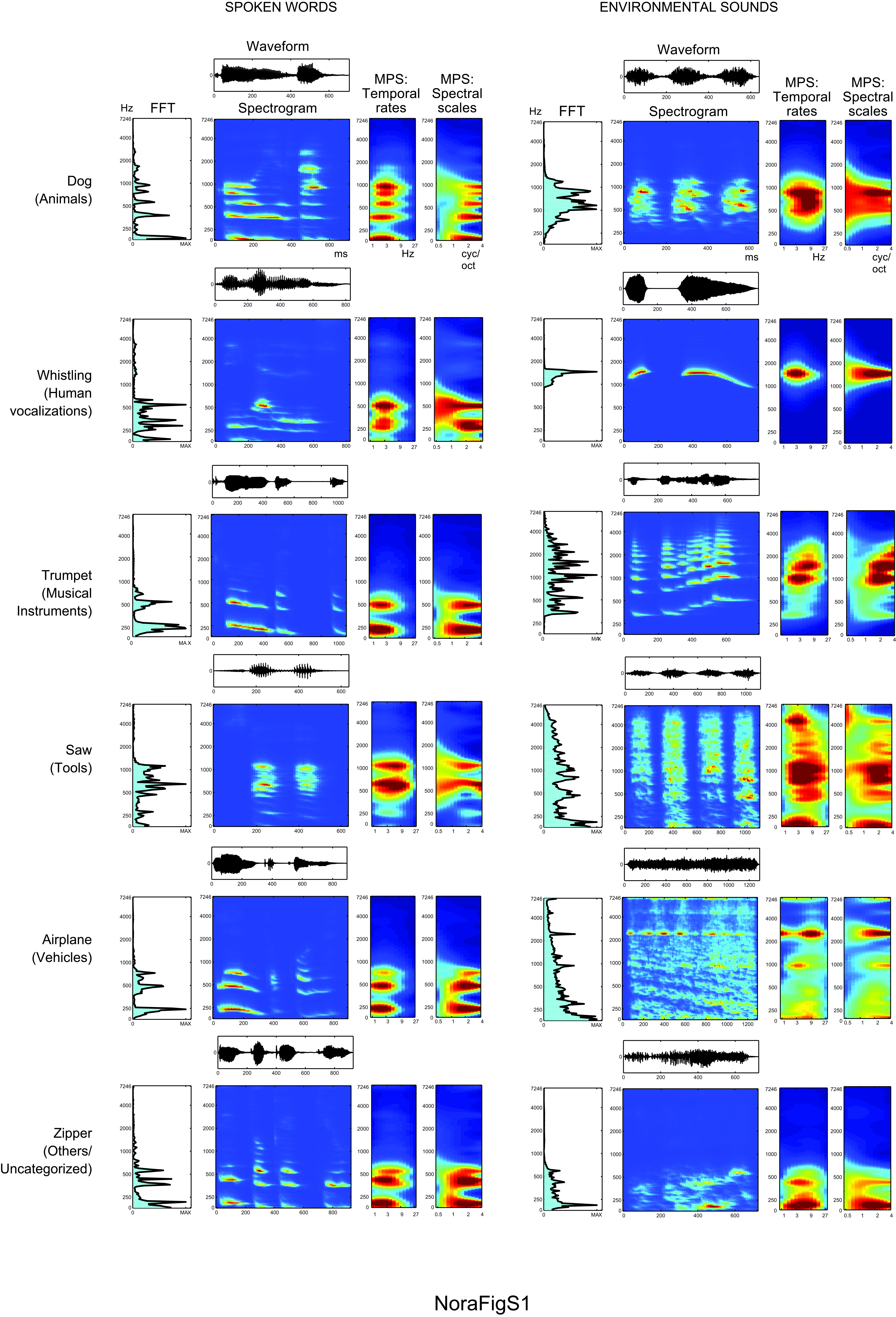

### Supplemental Figure 2

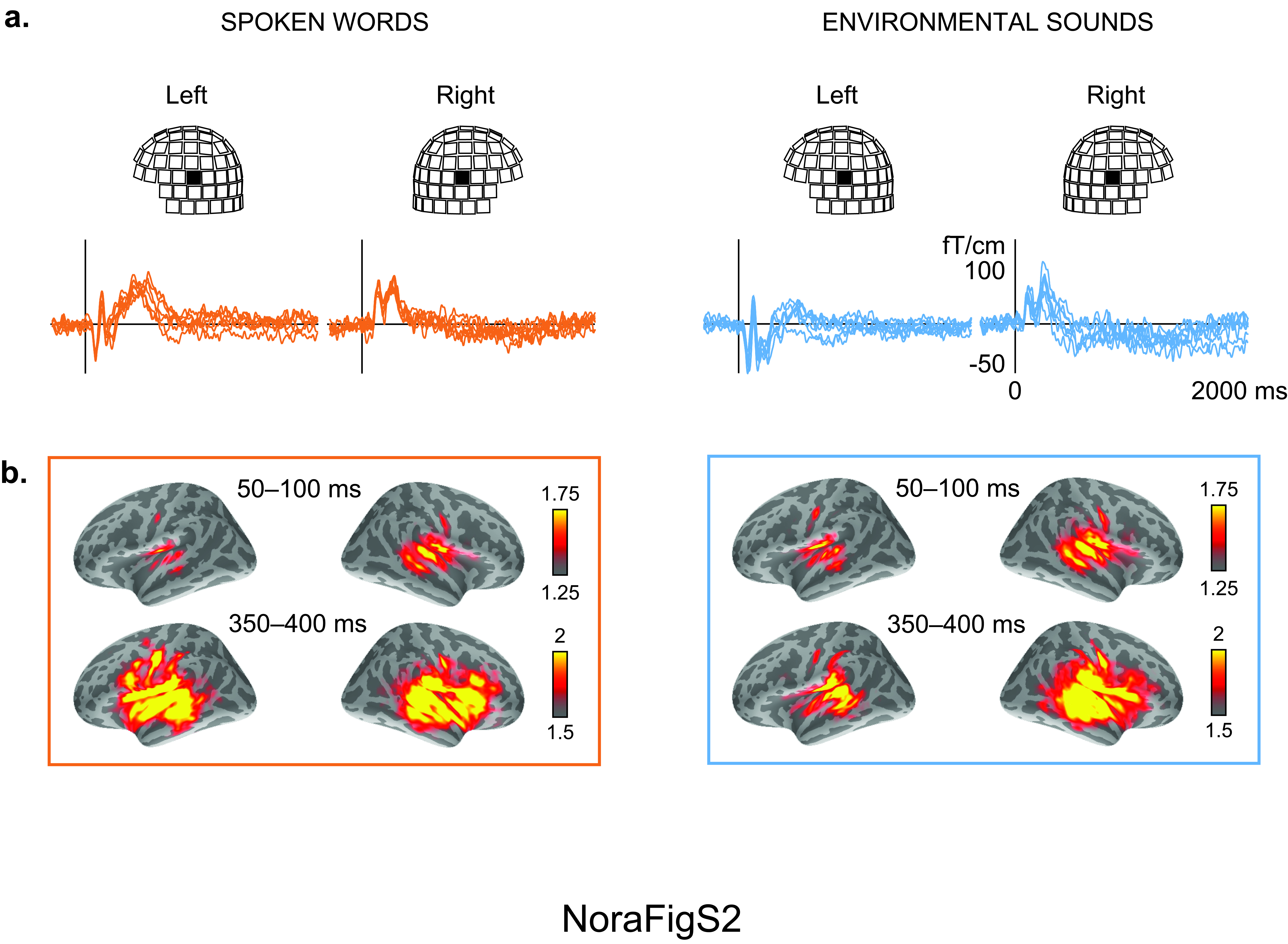

### Supplemental Figure 3

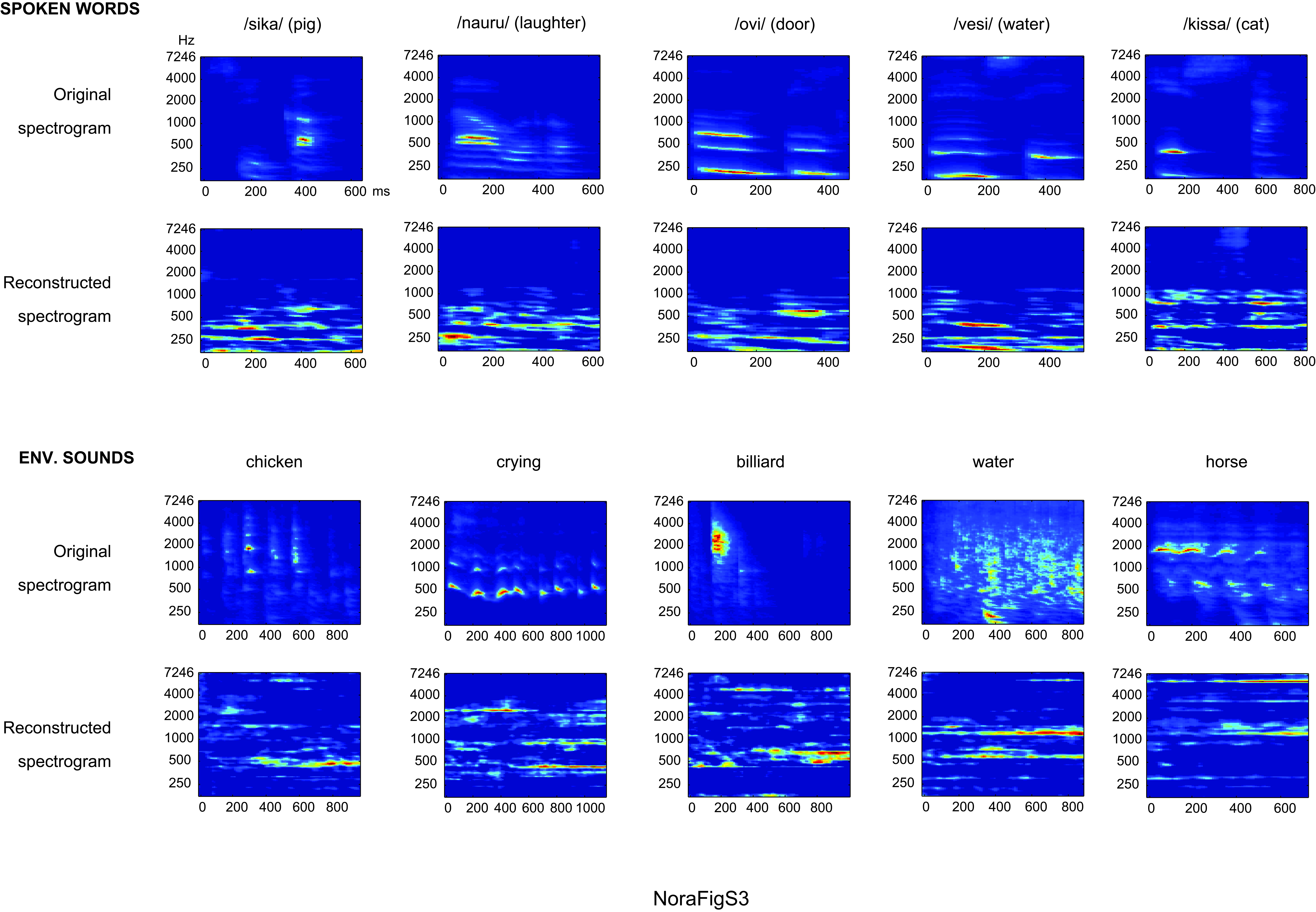

### Supplemental Figure 4

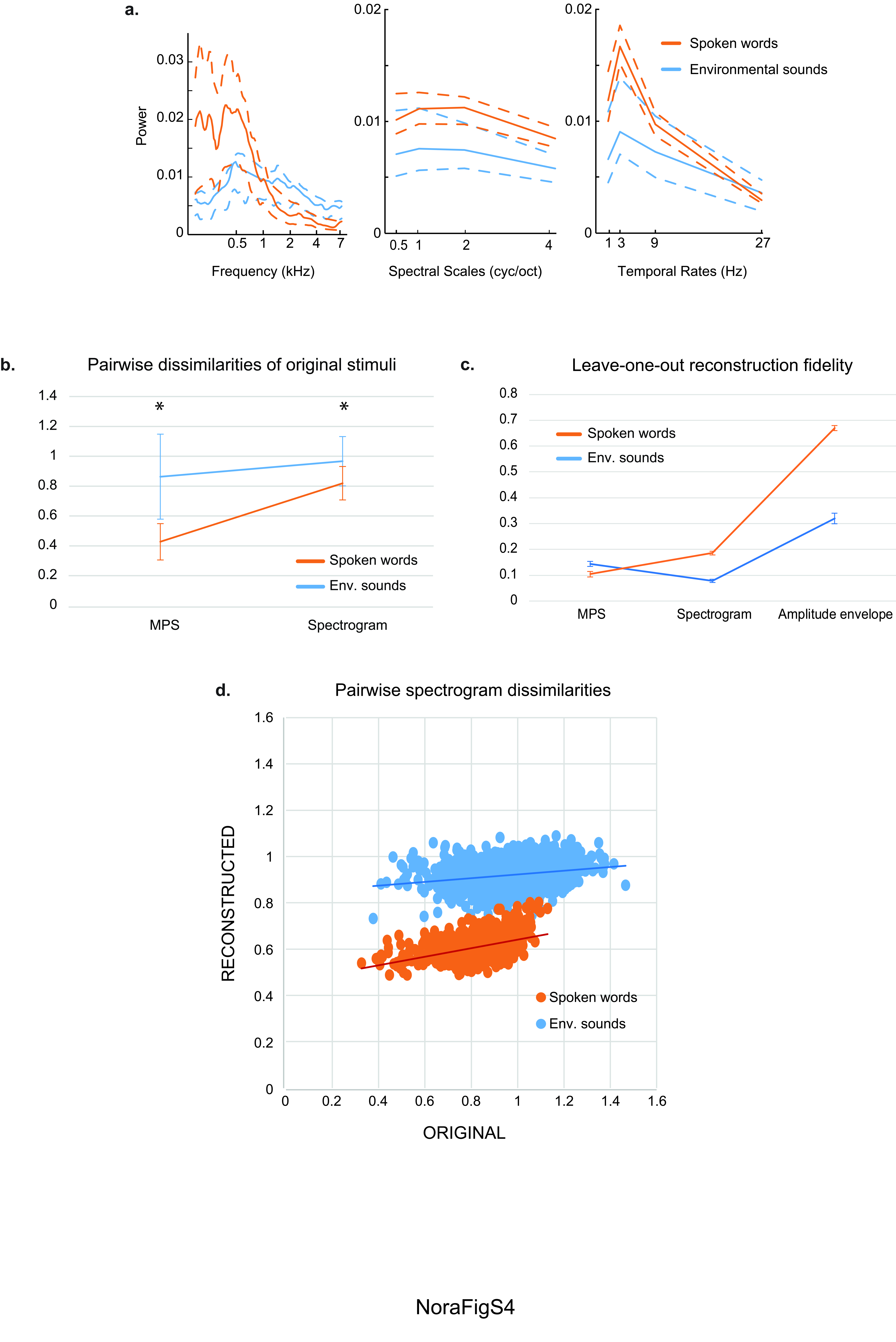

### Supplemental Figure 5

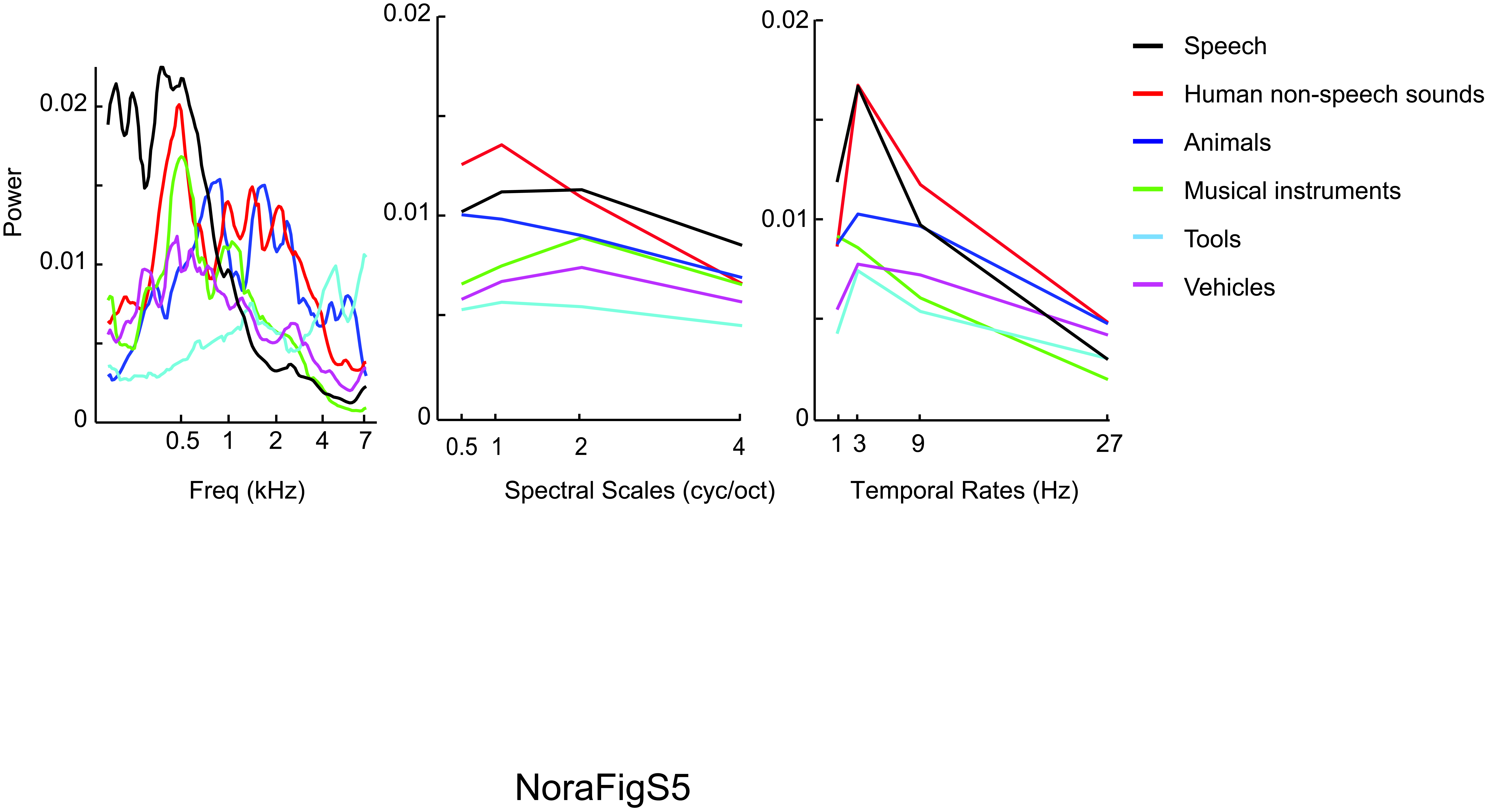

### Supplemental Figure 6

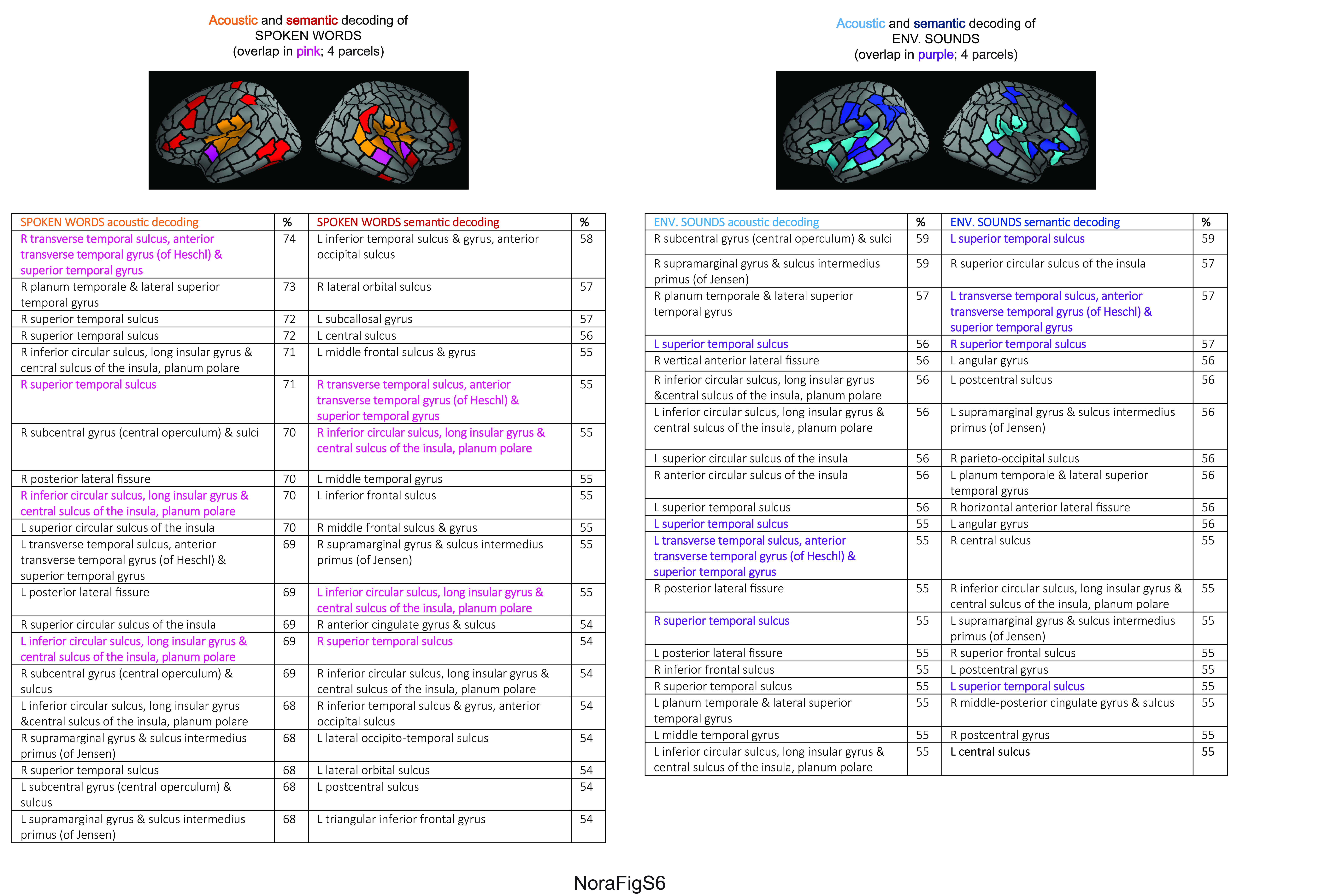
