## Supplemental Table 1 for "Dynamic time-locking mechanism in the cortical representation of spoken words"

Supplementary Table 1. List of stimuli: All items had one spoken word and one environmental sound exemplar.

| CATEGORY | ITEM | SPOKEN WORD (Finnish) | ENVIRONMENTAL SOUND |
|---|---|---|---|
| ANIMAL | Horse | hevonen | [horse neigh] |
| | Chicken | kana | [chicken bwaak-bwak-bwak] |
| | Cat | kissa | [cat meow] |
| | Dog | koira | [dog bark-bark] |
| | Sheep | lammas | [sheep baa] |
| | Cow | lehmä | [cow moo] |
| | Bird | lintu | [little songbird chirping] |
| | Pig | sika | [pig oink-oink] |
| HUMAN | Sneeze | aivastus | [female sneezing achoo] |
| SOUND | Yawn | haukotus | [male yawning] |
| | Crying | itku | [female crying] |
| | Laughing | nauru | [female laughing] |
| | Vomiting | oksennus | [male (mimicking) vomiting] |
| | Burping | röyhtäys | [male burping] |
| | Whistling | vihellys | [male whistling] |
| | Coughing | yskä | [male coughing] |
| | Accordion | haitari | [accordion chord] |
| MUSICAL | Flute | huilu | [three descending flute notes] |
| INSTRUMENT | Guitar | kitara | [two chords on guitar] |
| | Piano | piano | [two chords on piano] |
| | Drums | rummut | [one cycle of drum beat] |
| | Trumpet | trumpetti | [ascending trumpet tune with six notes] |
| | Organ | urut | [organ chord] |
| | Violin | viulu | [violin note with bow-change] |
| | Chainsaw | moottorisaha | [chainsaw running] |
| TOOL | Drill | porakone | [drill running on air] |
| | Saw | saha | [three draws of a hand-held saw] |
| | Match | tulitikku | [lighting a match] |
| | Hammer | vasara | [two strikes of a hammer] |
| | Knife | veitsi | [sound of sharpening a knife] |
| | Ambulance | ambulanssi | [ambulance siren] |
| VEHICLE | Car | auto | [car starting] |
| | Helicopter | helikopteri | [helicopter circulating] |
| | Train | juna | [train hitting the tracks] |
| | Ship | laiva | [ship foghorn] |
| | Airplane | lentokone | [airplane landing] |
| | Door | ovi | [door closing heavily] |
| OTHER | Doorbell | ovikello | [doorbell ding-dong] |
| | Telephone | puhelin | [a traditional telephone ring] |
| | Thunder | ukkonen | [thunder rolling afar] |
| | Water | vesi | [stream of water] |
| | Zipper | vetoketju | [zipper closing] |
| | Billiard | biljardi | [billiard balls hitting each other] |
| | Camera | kamera | [traditional camera shutter sound] |
